# Cumulative effects of nylon microplastic fibres and warming temperature on the behavior and physiology of marine threespine stickleback (*Gasterosteus aculeatus*)

**DOI:** 10.64898/2026.08.05.742354

**Authors:** Angelina L. Hajji, Kelsey N. Lucas

## Abstract

Populations are exposed to multiple anthropogenic stressors simultaneously; however, the combined effects are poorly understood. Climate change is starkly impacting marine ecosystems and consequently fishes, with warming temperatures and increases in frequencies and durations of extreme climate events. Concurrently, plastics, such as nylon used in fishing industries, are contaminating marine waters at unprecedented levels, with detrimental effects on fishes. Here we studied the cumulative effects of warming and nylon microplastic fibres on the behavior and physiology of threespine stickleback (*Gasterosteus aculeatus*) by exposing fish to conditions of 15°C and 20°C and nylon concentrations of 0, 1, 10, and 100 mg/g (mg nylon/g food) for 4 weeks. Feeding rates responded complexly to multiple stressors, as increasing concentrations of plastic reduced feeding rates, with warming having an antagonistic effect. Furthermore, we observed “coughing” behaviors in response to ingestion of microfibres and a unique reselection tendency of food items previously selected by conspecifics. Under warming conditions, critical thermal maximum (CT_max_) increased; however, exposure to plastics led to reductions in CT_max_ and thermal safety margins. Given these results, we anticipate reduced acclimation capacities, greater anxiety, and reductions in foraging efficiencies with increasing concentrations of plastic. Cumulatively, these stressors will yield greater energetic trade-offs and decreased accuracy in food selection with stark implications for marine ecosystem dynamics.

## INTRODUCTION

Animals are expected to encounter multiple stressors simultaneously in natural habitats; however, much of literature focuses on a single changing variable (Cheung et al., 2021; Guderley, 2004; Possatto et al., 2011). Our established literature base, hence, may profoundly underestimate the effects of various stressors, particularly as energetic budgets of an animal may shift under combined effects (Crain et al., 2008). Climate change is driving significant warming across global marine ecosystems with a three-fold increase in marine heat wave intensity since 1940, consequently impeding marine animals ( IPCC 2023; Lee et al., 2022; Marcos et al., 2025; Out et al., 2013). Concurrently, pollutants, especially plastics, are contaminating marine waters at unprecedented levels, leading to detrimental physical and cellular effects on marine animals (Law 2017). Marine fish populations – of great economic and ecological value – interact directly with the water around their bodies and are exposed to multiple anthropogenic stressors simultaneously. However, the cumulative effects of these remain poorly understood.

Many human stressors, such as pollutants or temperature changes, have *individually* been shown to have a variety of effects on fishes. These effects have been observed across all levels of biological organization and overarching ecological systems (reviewed below and in Hajji & Lucas, 2024). Exposure to warming temperatures past an individual’s optimum causes reduction in growth (Spinks et al., 2019), increases in physiological stress and decreasing metabolic performance (da Costa Barroso et al., 2020; Stewart et al., 2019), and altered behavioral responses such as increased aggression and indecision (Colby et al., 2022; Hajji & Lucas, 2026). Exposure to pollutants, including microplastic, has been shown to induce reproductive effects including tissue abnormalities and reductions or regressions in maturity (Hajji & Lucas, 2024; Maunder et al., 2007; Salierno & Kane, 2009); behavioral effects and disorders, sometimes through penetrating the blood-brain barrier (neurological effects) (Mattsson et al., 2017); and physiological and metabolic effects, such as gut dysbiosis and lipid metabolism disorders (e.g., excessive fat build up) that cause mortalities (Chen et al., 2020; Eroglu et al., 2015). We note that nylon is a dominating plastic source used in the fishing industry which contributes to a substantial amount of oceanic pollution. This plastic has received little attention in literature (Hajji & Lucas, 2024).

The *cumulative* effects of these stressors are widely unexplored but may be drastically compounding. Interactions between multiple stressors may be additive, synergistic, or antagonistic. Additive effects occur when the effect of each individual stressor is equal to their sum (Cabral et al., 2019). Synergistic effects describe a multiplicative effect, where the combined effect of stressors exceeds the sum of their individual effects (Cabral et al., 2019; Folt et al., 1999). Antagonistic effects are those where stressors interact in opposing manners where one stressor offsets (“cancels out”) the effect of the other (Cabral et al., 2019; Folt et al., 1999). Despite growing concerns with warming sea temperatures (Brierley & Kingsford, 2009; Harley et al., 2006; Poloczanska et al., 2016) and over 5 trillion particles of plastic in the ocean (Eriksen et al., 2014), limited literature has considered the cumulative effects of these stressors on fish. No known literature to date has focused on the cumulative impacts of nylon microplastics fibres and rising temperatures on fish, thermal tolerance, or behavior. To validate this, we performed a Google Scholar search on June 29^th^, 2026 which returned no relevant results (screened 10 pages with search terms: nylon, microplastic, warming, temperature, CTmax, thermal tolerance, behavior). Indubitably, understanding the combination of effects allows us to better understand interactions in natural habitats and responses of fish in wild ecosystems.

The combined impacts of temperature and plastic can be measured across levels of biological organization using behavioral, physiological, and growth endpoints. When used together, these endpoints aid in understanding how fishes cope with stressors, their potential reductions to fitness, and they allow us to infer changes to their energy budgets and allostatic load to expand the stressors’ mode of action from molecular to ecological. For instance, greater anxiety-associated behaviors or acclimation capacity (due to molecular upregulation) may exploit greater energetic reserves that decrease expenditure available for reproductive growth (e.g., gametes), with cascading population and ecological consequences. Using this adverse outcome pathway approach (see Ankley et al., 2010) and considering various physiological axes are highly beneficial in wholistically understanding stressors.

Behavioral endpoints are inherently driven by genetic and physiological interactions, are linked to damages in the nervous and endocrine systems, and significant ecologically relevant translations (Correia et al., 2007; Weis et al., 2001). Plastics are frequently associated with decreased satiety in animals (Hajji & Lucas, 2024), while elevated temperature often has opposing effects driving faster metabolic rates (Nunes et al., 2021). Literature however, remains limited on how particles affect individual and population feeding behavior, and the interactive effects of warming temperature (Wen et al., 2018). Behavioral assays, such as the black-white test (BWT), examine anxiety and exploratory responses of fish by observing their preference for darker environments over exposed areas of an arena (Bano et al., 2018; Blaser & Rosemberg, 2012; Norton & Gutiérrez, 2019). These behavioral endpoints provide key insight to the state of a fish and their perception of risk – potentially that may be associated with elevated temperature or plastic – in their environment.

For physiological endpoints, the Critical Thermal Maximum (CT_max_) is an important upper thermal tolerance limit with ecologically significant translations. The CT_max_ is the temperature at which physiological disorganization occurs, performance significantly declines, and, in fish, represents where death would occur in natural settings due to an inability to capture prey or evade predators (Beitinger et al., 2000; Morgan et al., 2018; Y. Zhang & Kieffer, 2014). CT_max_ values vary between individuals, populations, and species, and aid in understanding thermal performance, where ambient temperatures near CT_max_ values would indicate declines in performance (McKenzie et al., 2021). Established CT_max_ values, noted as a loss of equilibrium and dorso-ventral positioning in fish, can additionally be used in determining Acclimation Response Ratios (ARRs) and Thermal Safety Margins (TSM). ARRs are ratios between CT_max_ values at different ambient temperatures which are used to determine whether shifts in upper thermal tolerances are equivalent to experienced change in ambient temperature (Morley et al., 2019).

ARR values less than one indicate decreased acclimation capacity (Morley et al., 2019). TSM, calculated as the difference in ambient temperature from the CT_max_, predict the tolerable temperature buffer region before loss of function (Sunday et al., 2014). The ARR and TSM are especially useful in identifying individual and population acclimation capacity and vulnerability. While previous research has demonstrated reduced acclimation capacities at warming temperatures in many species (De Bonville et al., 2025; Ruthsatz et al., 2024), no known literature has investigated the impacts of plastic particles on thermal tolerance, despite their documented impacts on gill filaments and energy budgets (Cao et al., 2023; Yin et al., 2018).

To better characterize mechanisms and axes of impact, particularly in regard to energetic budgets, combining growth endpoints such as changes in weight, length, or condition factor with the previous endpoints is useful. Growth is contingent on energy available after fulfilment of necessary costs such as digestion, cellular repair, and activity (Neubauer & Andersen, 2019; Volkoff & Rønnestad, 2020). Both plastics and temperature have individually been found to decrease overall growth due to increasing allostatic energy demands leading to reduced fitness overall (Alfonso et al., 2021; Liang et al., 2023; W. Wang et al., 2020), and we would expect their combination to amplify these effects. Combining these growth, physiological, and behavioral axes allows for a more wholistic understanding of overall animal well and fitness and for studies to better predict ecological effects.

Threespine stickleback (*Gasterosteus aculeatus*) are of great ecological importance as a prey species and are globally distributed, making them a useful model organism for testing human stressors on marine ecosystems (reviewed in Hajji & Lucas, 2026). Threespine stickleback in the Canadian North Pacific are considered genetically conserved across many natural environmental gradients such as salinity (freshwater to marine), depth (0-100 m), and temperature (4°C-20°C), with well documented physiological responses (Froese and Pauly 2022). As with other fishes, stickleback have demonstrated an ability to acclimate to many natural stressors, with most changes attributed to their phenotypic plasticity (Day & McPhail, 1996; Mazzarella et al., 2015; Mottola et al., 2022). As such, many genomic, physiological, and ecological connections have been previously established making them an ideal organism to study the effects of anthropogenic – pollutant and temperature – stressors on marine animals, particularly under shorter timescales. Populations in the Northern Pacific (surrounding Vancouver Island coast) face local warming and extreme climate events (e.g., recent heat waves) and plastic concentrations upwards of 600 particles per cubic meter (Collicutt et al., 2019). Fish populations inhabiting these waters are observably contaminated, and as such this study system is effective in understanding and predicting the impacts and risks of these stressors across local and global marine ecosystems.

Using the stickleback as a model, we investigate 1) effects of plastic exposure on fish function, 2) effects of multiple stressors – plastics and warming temperatures – on fish (antagonistic, synergistic, or additive), and 3) thermal capacities of fish under stressed conditions. We hypothesize that stressors will adversely affect fish, will combine in an additive or multiplicative manner, and specifically will reduce the thermal capacity of the fish, as warming temperatures and microplastics will impair functionality and increase the proportion of a fish’s energy budget that is expended for survival.

## METHODS

### Ethics

All work was approved by the Bamfield Marine Science Centre Animal Care Committee and the University of Calgary Life and Environmental Sciences Animal Care Committee, performed under the Animal Care Protocols RS-24-01 and AC24-0055, respectively.

### Preparation of Microplastic Food Samples

Commercially available monofilament nylon fishing wire was purchased from Mckanti. This nylon sample was validated for its chemical structure (i.e., verifying plastic was chemically monofilament nylon 6-6, without additives) using Fourier transform infrared spectroscopy (FTIR) at the University of Calgary Faculty of Science Instrumentation Facility (Figure 1a). The wire was prepared into a homogenous mixture of microplastics <5 mm (de Souza Machado et al., 2018; Graham & Thompson, 2009; Gray & Weinstein, 2017; Grigorakis et al., 2017; Jovanović et al., 2018; Kwak & An, 2020). Initially, the nylon wires were cut using scissors into 5 cm strands and then placed in a coffee grinder (Spectrum Brands Inc Model CBG110) for shredding in interval durations between 2-5 minutes. Then strands were transferred into a food processer (Cuisinart^®^ Model DLC-2AC) to further decrease sizing, for durations up to about three minutes. Remaining strands identifiably longer than 5 mm were manually cut. The mixture of microplastics was then sieved twice using a 5 mm sieve and a 1 μm sieve, eliminating plastics larger and shorter respectively, maintaining the microplastic size class. This homogenous mixture of microplastics was then added to the food samples prepared daily in the respective quantities for control, low, medium, and high plastic exposure treatments (0 mg/1 g, 1 mg/1 g, 10 mg/1 g, 100 mg/1 g; mg microplastics / mg feed; see Table 1 and Supplementary Table 1). These quantities reflect environmentally relevant concentrations (Sun et al., 2021).

**Figure 1.**
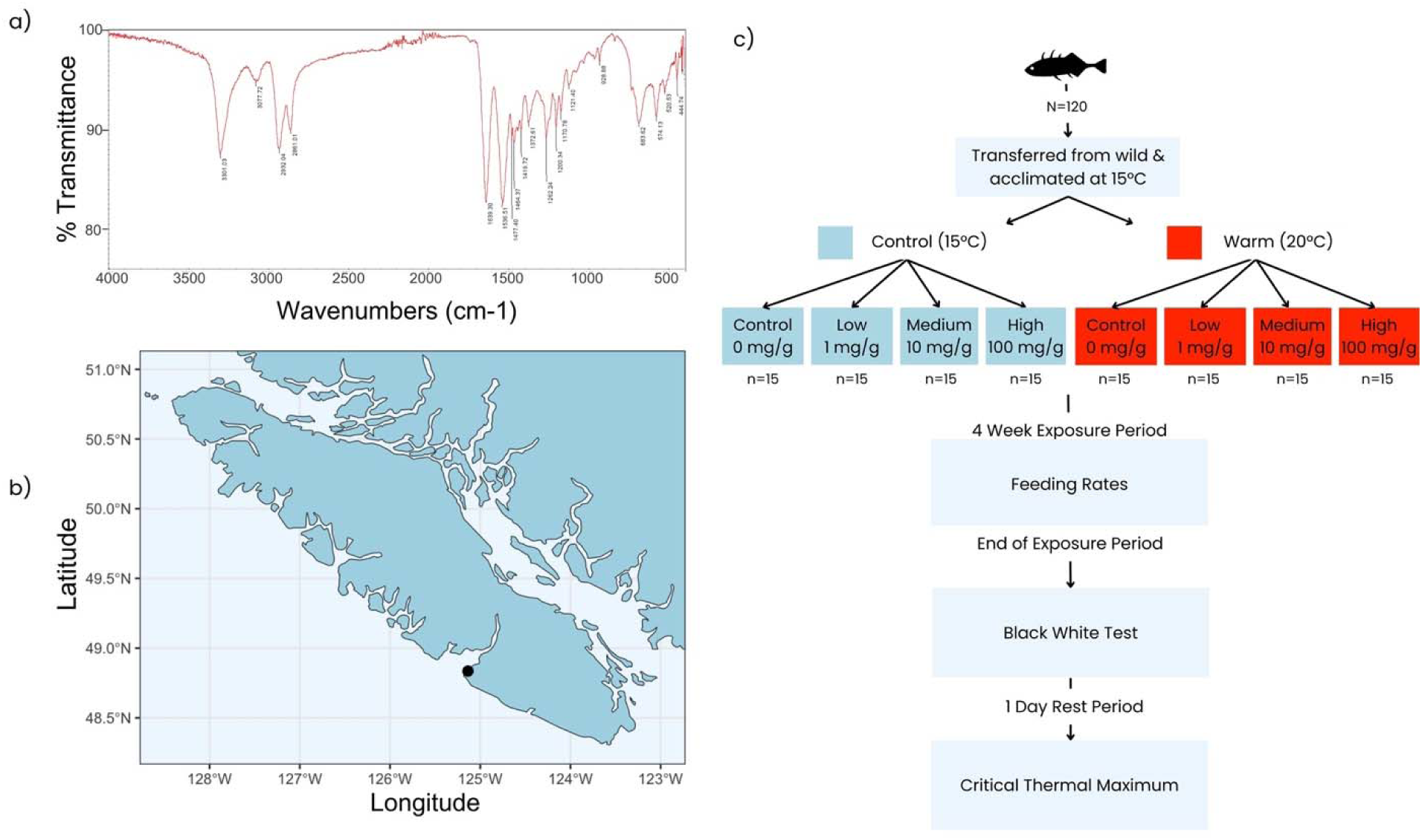
a) FTIR Analysis conducted validating match for Nylon (Polyamide 6 + Polyamide 6,6). b) Collection site for threespine stickleback adjacent to the Bamfield Marine Science Centre in Canadian Pacific Management Area 23 (48.8141056° N, -125.1565139° W). c) Overview of experimental design. Marine threespine stickleback (N = 120) were collected using minnow nets and randomly sorted into one of eight treatments (3 replicates tanks for each; n = 5), within a fully crossed experimental design.

**Table 1.** Tank temperature values (°C) and plastic concentration exposures (mg nylon / g bloodworm) for treatments during four-week exposure period.

| Treatment | Tank Temperature (°C) | SE Temperature | Microplastic Feed Concentration (mg nylon / g bloodworm) |
| --- | --- | --- | --- |
| Control | 14.2 | 0.11 | 0 |
| Control Low | 14.15 | 0.11 | 1.05 |
| Control Medium | 15.37 | 0.79 | 10.09 |
| Control High | 14.3 | 0.11 | 108.31 |
| Warm | 19.73 | 0.33 | 0 |
| Warm Low | 19.88 | 0.31 | 1.03 |
| Warm Medium | 20.07 | 0.31 | 10.09 |
| Warm High | 20.16 | 0.31 | 100.07 |

### Fish Sampling and Husbandry

Adult threespine stickleback (N = 120) were collected using Gee minnow traps from a single site adjacent to the Bamfield Marine Science Centre (Canadian Pacific Management Area 23; 48.8141056° N, -125.1565139° W; Figure 1b) and transferred to standard fish lab conditions at the station (∼15°C, 16D:8N). To maintain environmental water quality conditions upon transfer, water was fed directly from the ocean and filtered through a 1 μm filter to remove environmental particle matter, before entering the flow through system, where each tank was independently serviced. To prevent the entry of microplastics used during the experiment into natural waterbodies, the outflow of tanks was additionally suited with a 0.1 μm filter. Upon arrival at the lab, fish were randomly sorted into 24 tanks with five fish per tank (70 cm x 50 cm x 30 cm; see Figure 1c). Upon sorting fish were given two weeks to recover from any handling stress and to establish social hierarchies. Fish were fed standard bloodworm diet (1.5 g / tank) during this period.

### Exposure Design and Timeline

Prior to treatments, baseline behavioral assays were conducted (see *Behavioral Assays* below) for all individuals. These sticklebacks were then acclimated for four weeks for their respective treatments (three tank replicates per treatment) following a fully crossed experimental design across 1) temperature: control (15°C) and warm (20°C) temperature treatments; and 2) plastic exposure gradients: control (0 mg/g), low (1 mg/g), medium (10 mg/g), and high (100 mg/g) (refer to Figure 1c for experimental set up and Table 1 for tank data). Tanks were allotted 0.3 g bloodworms per fish (1.5 g per tank; >10% body weight) once daily accordingly to their treatment (see *Preparation of Microplastic Food Samples*) following Hajji & Lucas (2026). The amount provisioned was above maintenance proportions to ensure food was not a limiting factor for health in this experiment. Tank temperatures were controlled using submersible Fluval^®^ heaters, with the warming temperature increased to its target temperature within 24 h. Following the treatment period for the experiments, growth, behavioral, and CT_max_ measurements were performed.

### Behavioral Assays

The behavioral responses of fish to the treatments (N = 120) were determined through general observations and two behavioral assays including 1) feeding behavior test and 2) the black-white preference test (BWT). For the feeding behavior test, the fish were provisioned their daily allotted feed for their respective treatment and the total duration for food consumption (per tank) were observed and recorded. The feeding rate was then calculated by tank as:

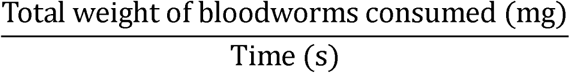

For each tank, 10 feeding behavior tests were collected for each tank throughout the exposure period.

The BWT was performed before and after the four-week acclimation period for all treatments and recordings were done with GoPro HERO 10s (60-240 fps, linear mode) following methods from Hajji & Lucas (2026) and Norton & Gutiérrez (2019). Briefly, in the BWT, an individual was individually transferred into a transparent tank (33.5 cm x 17 cm x 22 cm) with black and white opaque covers that divide the tank into a white zone and a black zone. The individual’s behavior was recorded from above for a duration of 5 minutes. The behavior was manually quantified from the footage by recording the time spent in the white zone and the number of crosses made from the black to the white zone. The researcher was blinded with respect to the treatment during these analyses. For all assays, the water in the assay tanks were changed after every second fish tested and between tank treatments; temperature in the test tank was maintained at the temperature the fish were acclimated to.

### General Health Monitoring and Growth

Growth metrics including weight and length (including caudal fin) were recorded for each fish (N = 120) before and after the four-week exposure period. These data were used to calculate the condition factor (K) using the formula provided by Paul 1983:

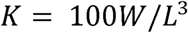

Where K is the condition factor, W is the weight of the fish in grams, and L is the length of the fish in cm. An additional exponent value of 3.03 (*L*^3.03^) was used to calculated following Hajji and Lucas (2026) and data from fishbase.org, which more specifically captures threespine stickleback growth.

As many pollutants have been found to have off-target effects than the research focus, we included general health monitoring throughout the duration of exposure. Monitoring followed a qualitative approach, where observations of fish (morphology or behaviors) in standard housing arena were logged daily. Observations such as conspecific interactions or individual feeding were recorded as well as descriptions of the feeding process and interactions with microplastics. In addition, recording of any abnormal health endpoints (e.g., general lethargy) and morphological observations were included. Two fish from a single tank (warm, high plastic treatment) were euthanized due to detection of humane endpoints (see Results: *General Health Monitoring and Growth*) during the final day of the experiment, following all behavioral assays but prior to the CT_max_ trial.

### Critical Thermal Maximum (CT_max_)

Following the four-week exposure period, fish were provided a single day recovery period between the BWT test and the CT_max_ trials; fish were fasted for 24 h during this period. For these, four of five fish from each tank were selected randomly and placed individually in 1000 mL glass jars in an experiment tank and provided a 15-minute acclimation period prior to the trial commencing. For the warm high plastic treatment 11 fish were sampled (n = 4, 4, and 3), due to fish condition (see Results: *General Health Monitoring and Growth*), resulting in a total N = 95. CT_max_ measurements typically show little variation among individuals within treatments, and as such there is no anticipated impact of the uneven sample size on the results (Becker & Genoway, 1979; Raby et al., 2025). The CT_max_ trials began at the temperature the fish were acclimated to, to avoid temperature shocking the fish prior to the acute trial test (e.g., 20°C treatment began trial at approximately 20°C). Methods for CT_max_ followed Hajji and Lucas (2026, see Supplementary Table S1 therein), where the temperature was raised approximately 0.33°C min^-1^ until the fish lost equilibrium noted as the point where it can no longer maintain dorsoventral positioning (Becker and Genoway 1979; Beitinger et al., 2000). This is the point where fish would not survive in natural conditions (e.g., loss of predation escape). To calculate TSM the following formula was used (Morley et al., 2019):

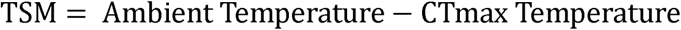

The ARR was calculated following Claussen (1977):

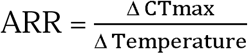

where lower ARR values represent lower acclimation capacity.

### Statistical Analysis

Data were analysed using R (2023.03.1+446) (R Core Team, 2024). Data analysis used a combination of 1) linear mixed effects models (LMER) (lmerTest_ 3.1.3), 2) general additive models (GAM) (mgcv_ 1.9.1), and 3) odds ratios (OR) using general linear mixed effects (GLMER) (lme4_1.1.35.5) as appropriate to characterize combined effects (Bates et al., 2015; Harrison et al., 2018; Kuznetsova et al., 2017; Wood, 2017). All random effects were treated as random intercepts. Where applicable, QQ plots and Shapiro-Wilk’s tests were used to validate normality and Barlett’s tests for the assumption of homogeneity of variance. Where appropriate, backward selection was then conducted using the Akaike Information Criterion (AIC) (Zuur et al., 2009). Where significance (p < 0.05) existed in LMER and GAM analysis, pairwise comparisons were conducted with estimated marginal means (EMMs) with Sidak adjustment (emmeans_1.10.3) (Lenth, 2023).

More specifically, feeding rates were analyzed using LMER with plastic and temperature as fixed effects and tank and day (to account for temporal variation) as random effects. An initial Shapiro test on the feeding rate data yielded high W values, however residuals of an initial LMER invalidated normality assumptions. Feeding rates were log-transformed to account for this. The LMER was then refit on the transformed data. The final model following backward selection included temperature and plastic as independent fixed effects and day as a random effect.

BWT yielded two types of data, duration (i.e., time spent in the white zone) and cross (i.e., crosses to white zone) that were analyzed independently. Duration data were analyzed using LMERs; the selected model used plastic, temperature, and their interaction as fixed effects and tank as a random effect. Raw EMMs were required for pairwise comparisons. The cross data, as counts, were analyzed using a GAM with Poisson family distribution, REML method, and used plastic and temperature as independent fixed effects, with tank as a random factor.

Following initial analysis of all behavioral data (feeding rates and BWT counts), odds ratios (ORs) were computed (Szumilas, 2010; Tenny & Hoffman, 2023). ORs effectively predict how strongly an outcome is associated with a given exposure treatment (Szumilas, 2010). OR was calculated for three measured variables: feeding rates, duration, and cross. For each, the data were split by the global median value as OR calculations require categorical data. To do this, data were binomially divided into two groups: 1) values below median = 0, and 2) values above median = 1. The GLMER used in the feeding rates OR calculation included temperature and plastic as fixed effects and both day and tank as random intercepts. The GLMER for the BWT cross OR calculation included temperature as a fixed effect and tank as a random effect. The OR was then used in all cases to calculate the likelihood of an events occurrence, compared to the control (i.e., odds occurring in X treatment / odds occurring in control).

The selected LMER models for CT_max_ and growth data used temperature and plastic concentration as fixed effects and tank as a random effect, following methods outlined above.

## RESULTS

### Behavioral Assays

Feeding rates responded complexly to individual and combined treatment groups; all treatments differed significantly from the control (Figure 2; Supplementary Tables 2-4). Feeding rates decreased significantly with plastic concentration, with the lowest rate observed under the 100 mg/g plastic concentrations (Figure 2; Supplementary Tables 2-4). Warm temperature conditions significantly increased feeding rates compared to the control temperature treatment (Supplementary Tables 2-4) and antagonistically interacted to partially offset plastic concentrations. This effect was observed at all plastic exposure treatments (Figure 2; Supplementary Tables 2-4). Warming was 11.5 times more likely to experience faster feeding rates, compared to control (Supplementary Table 5). Microplastic exposure led to a greater likelihood of slower feeding. The 1.0, 10, and 100 mg/g plastic exposures were 3.7, 1058.1, and 10680.4 times as likely to experience slower feeding rates (i.e., below the median threshold; Supplementary Table 5).

**Figure 2.**
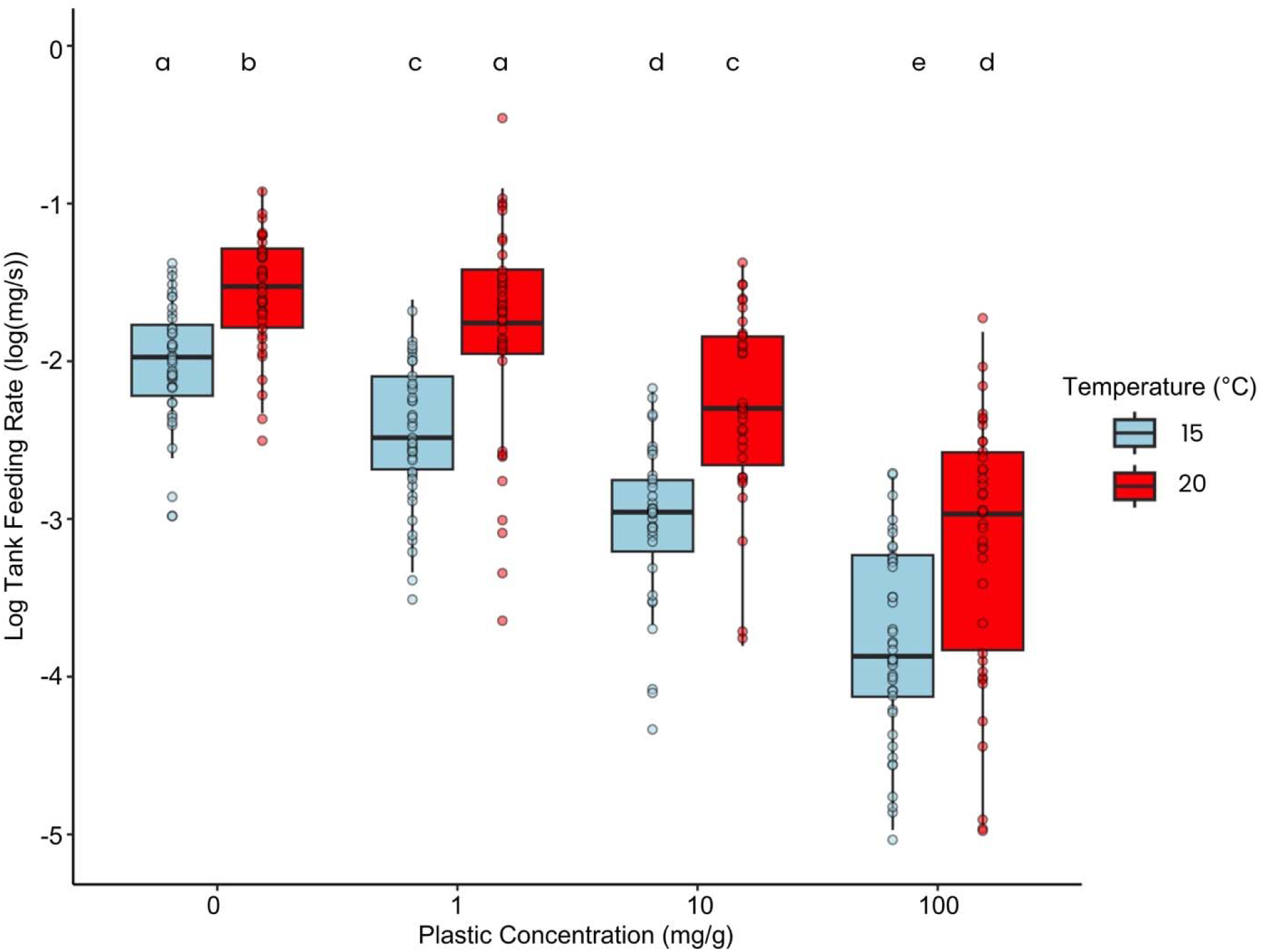
Tank feeding rate results (log(mg/s) per tank) across temperature (15°C and 20°C) and microplastic (0, 1, 10, 100 mg/g) treatment groups. Bars represent 1.5 x the interquartile range, dots represent individual data points (per tank per day) (n = 32-42). Letters indicate significance (see Supplementary Tables 2-4).

In the BWT, duration (time spent in the white zone) and the number of crosses were the highest in the warm treatment, with microplastic exposure having a significant impact on time spent in the white zone (Figure 3a; Supplementary Tables 6 and 7). Temperature was the only significant factor impacting crosses in the model (Supplementary Table 8). Warming temperature significantly increased the number of crosses from the black to white zone (Figure 3b; Supplementary Table 8). Fish were 1.86 times more likely to experience high cross counts in the warm treatment, in comparison to the control (Supplementary Table 9). Plastic treatments had an opposing effect; 1.0, 10, and 100 mg/g plastic treatments were respectively 10, 2.7, and 7.1 times less likely to experience high cross counts than the control (Supplementary Table 9).

**Figure 3.**
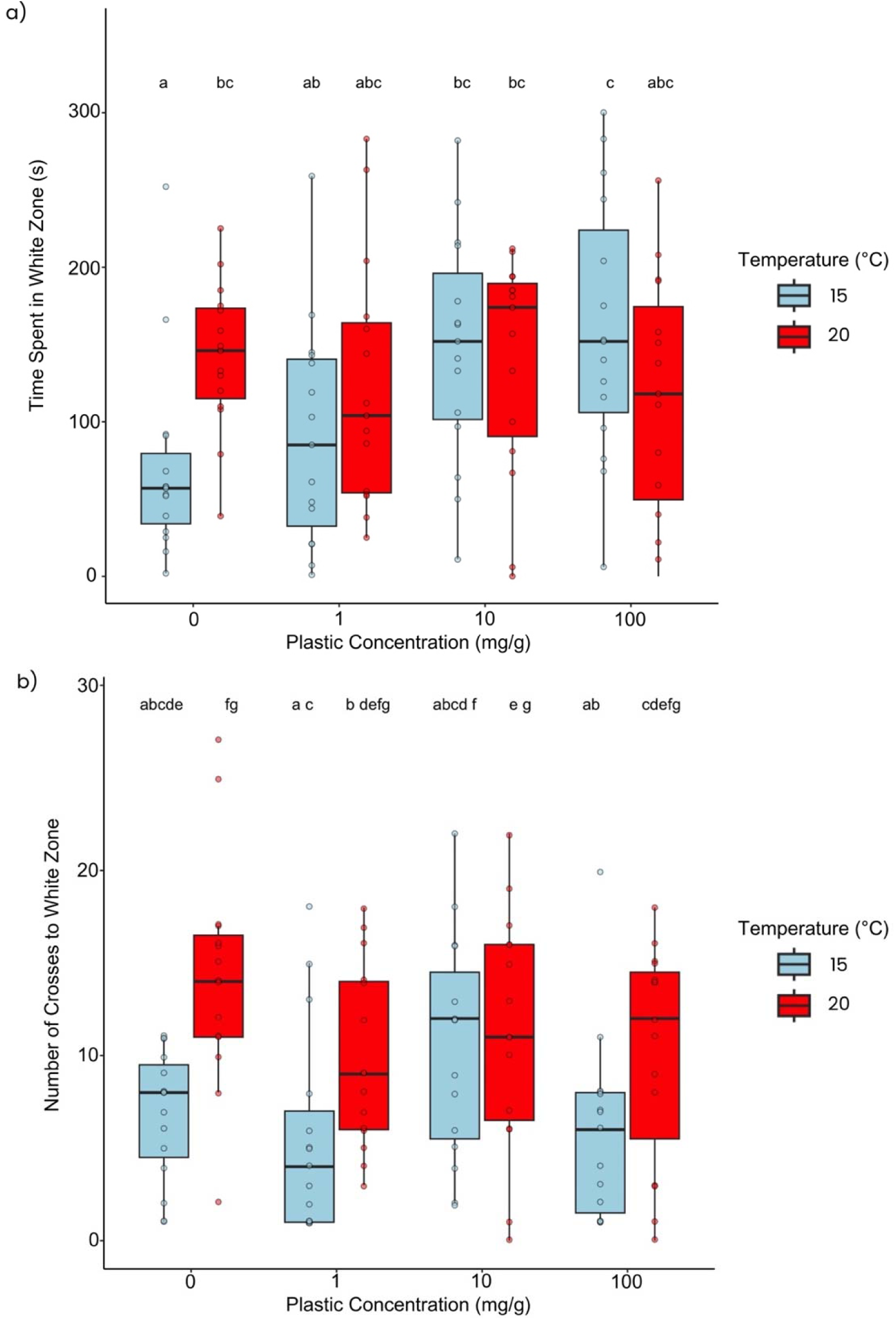
Black-White Test results for temperature (15°C and 20°C) and microplastic (0, 1, 10, 100 mg/g) treatment groups. a) Time (s) spent in the white zone of the tank and b) frequency of crosses to the white zone of the tank (n = 15). Letters indicate significance (Supplementary Tables 6-8).

### General Health Monitoring and Growth

Across all treatments, fish demonstrated no statistical differences in their change in length and weight (Figure 4a-b, Supplementary Tables 10-11). There was however a non-significant trend with plastic concentration treatments demonstrating higher weights despite similar lengths (Figure 4a-c) (Supplementary Tables 10-11). Further, there were no differences detected in condition factor among treatments following pairwise comparisons, despite significance detected from high plastic in the respective LMER (Figure 4c; Supplementary Tables 12-13).

**Figure 4.**
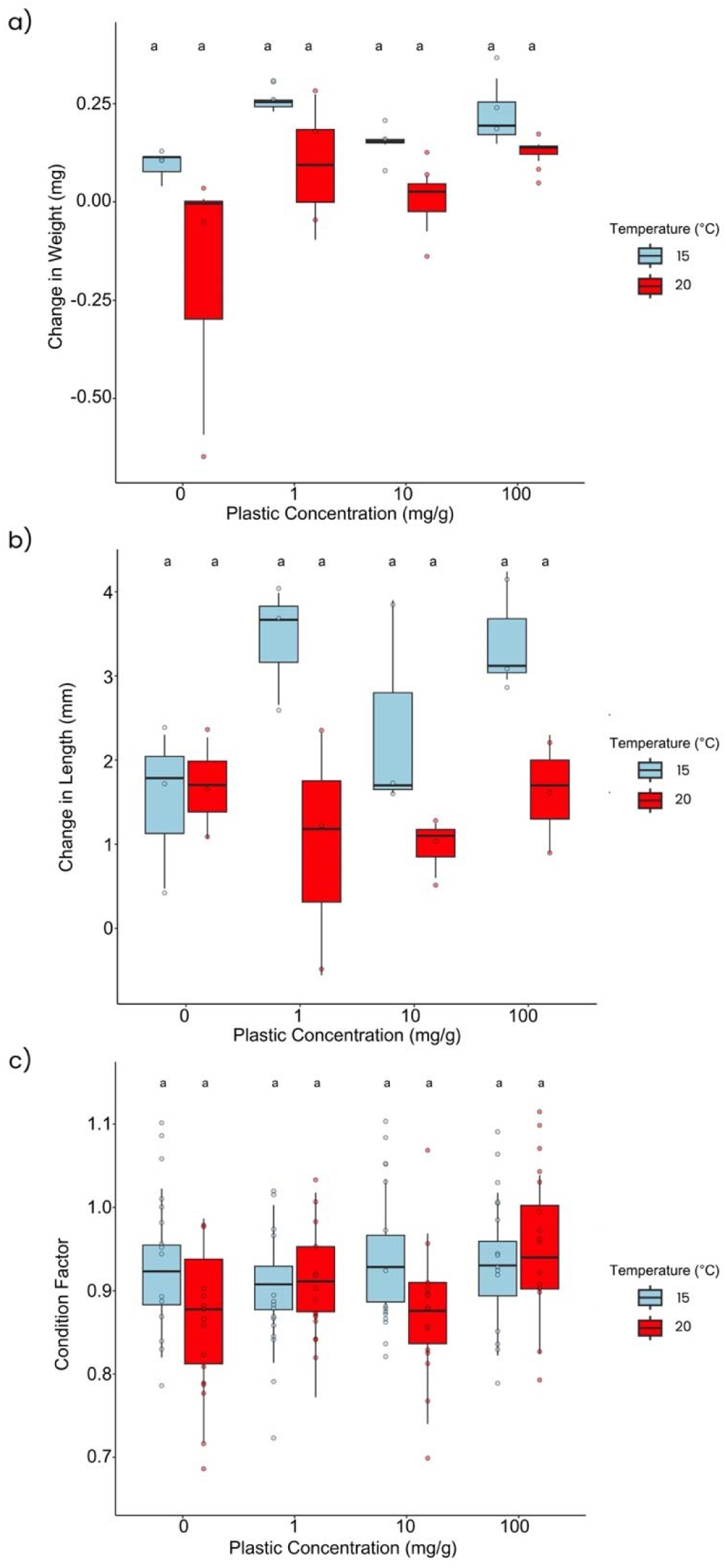
Boxplots of morphological changes observed in growth metrics during experiment, bars represent 1.5 x the interquartile range. a) Average change in weight (mg) by tank (n = 15), b) average change in length (mm) by tank (n = 15). Dots represent tank averages (n = 3). and c) change in condition factor by individual (n = 15); dots represent individuals. Letters indicate significance (see Supplementary Tables 10-13).

The general health monitoring logs captured qualitative impacts of both temperature and plastics on the fish. Notable aggressive behavior occurred in the warm treatments with higher instances of “tripoding” (lifting of spines in stickleback) in response to researcher presence. Biting or chasing intraspecific aggressive encounters were seemingly more prominent in warm treatments in comparison to the control. Furthermore, during temperature checks, attacks of the probe increased with temperature. This effect was seemingly counteracted by plastic concentration exposure. In the warm treatment, fish followed fingertips outside of the tank and approached with a tripoding response. There were greater instances of this behavior recorded in lower plastic exposure concentrations regardless of temperature. Fish displayed frequent hiding under objects, submissive backing when tank is approached, increased with increasing plastic concentrations, regardless of temperature. This however, appeared to be offset by temperature. Intraspecific aggression was noted for all treatments, however appeared greater in tanks with high plastic exposure as individuals often “fought” over bloodworms despite an abundance present. During feeding we observed a coughing behavior during some instances of microplastic ingestion (Figure 5a-b) and a unique conspecific reselection of food particles previously selected and rejected by first individual (Figure 5c). We further observed under many instances, false selection of nylon microplastic particles over available food particles (Figure 5d). In the 100 mg/g plastic treatments, individuals developed distended abdomens. Two fish from the warm high plastic treatment (from a single tank) were euthanized during the final week of the study, after behavioral assays were completed but before CT_max_ data were collected, due to persistent gut swelling observed, difficulty swimming, and an inability to maintain dorso-ventral positioning (see Figure 5e).

**Figure 5.**
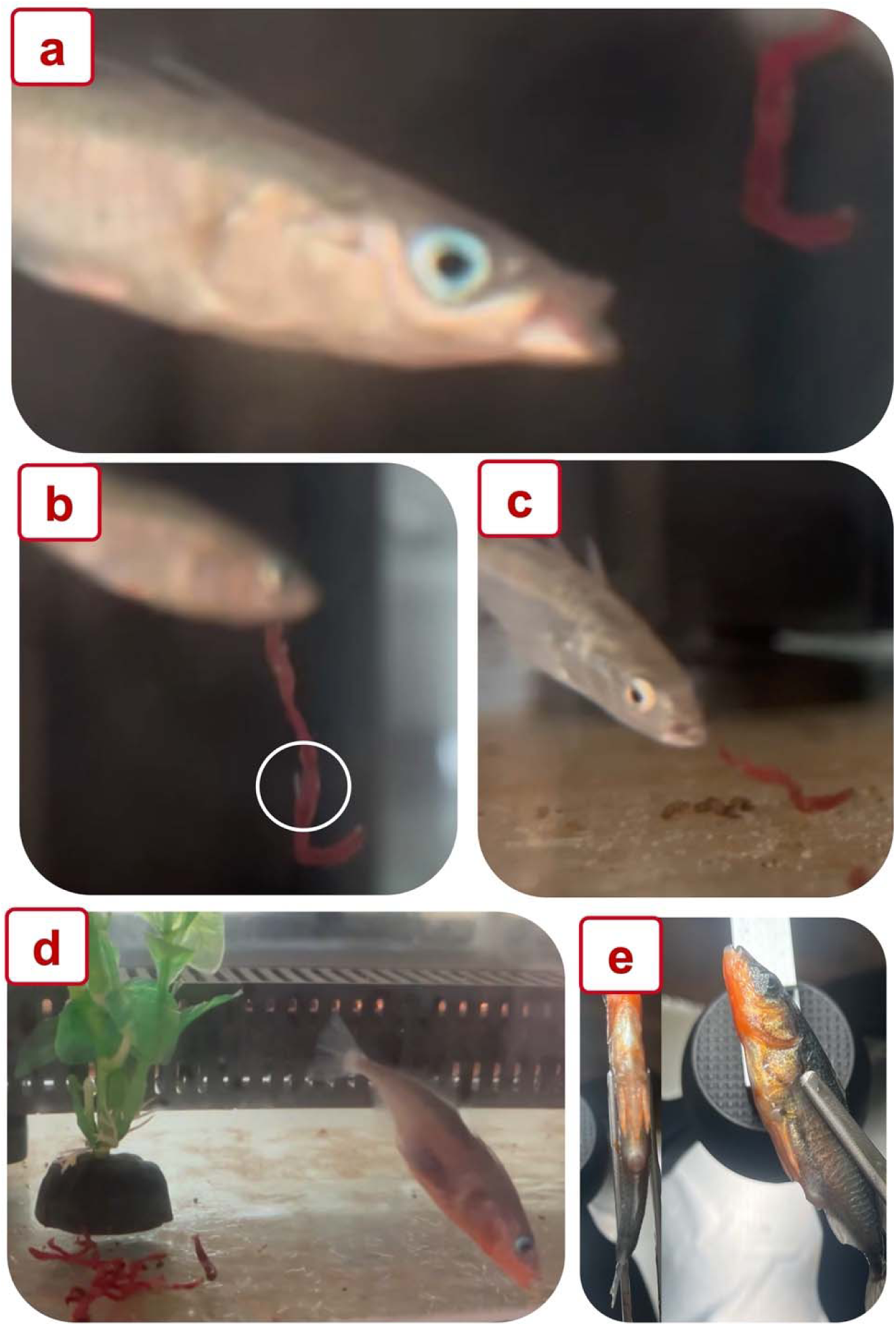
Images of behaviors and morphology exhibited by individuals during experiment. a) Example of “coughing” behavior in stickleback following microplastic ingestion and rejection. b) Image of microplastic in food particle rejected by individual in (a). c) Conspecific reselection of food particles previously selected and rejected by first individual observed in (a). d) False selection of plastics over food particles. e) Image of abdominal swelling in individual unable to right itself (100 mg/g + warm treatment).

### Critical Thermal Maximum (CT_max_)

Both temperature and plastic exposure significantly impacted CT_max_ values (Figure 6 and Supplementary Table 14). CT_max_ values were significantly higher in the warm treatment (34.5°C ± 0.03) in comparison to the cold (32.9°C ± 0.09) (Figure 6; Table 2; Supplementary Tables 14-16), with an ARR value of 0.36 (see Table 2). The TSM decreased from 18.70°C in the control to 14.77°C in the warm treatment (Table 2). Warming treatments had significantly higher CT_max_ values than their cold treatment counterparts, regardless of plastic exposure (Table 2; Supplementary Tables 14-16). Plastic exposure complexly impacted CT_max_ values, seemingly following a biphasic response with the medium levels of plastic concentration experiencing the lowest CT_max_ values for their respective temperature treatments (see Figure 6, Table 2, and Supplementary Tables 14-16). In the control temperature treatments, exposure to plastics significantly reduced CT_max_ values between 0.591.09°C, decreasing TSM values (Table 2). A similar trend was observed in the warm treatment (see Figure 6 and Table 2). The CT_max_, ARR, and TSM values for all treatments can be found in Table 2.

**Figure 6.**
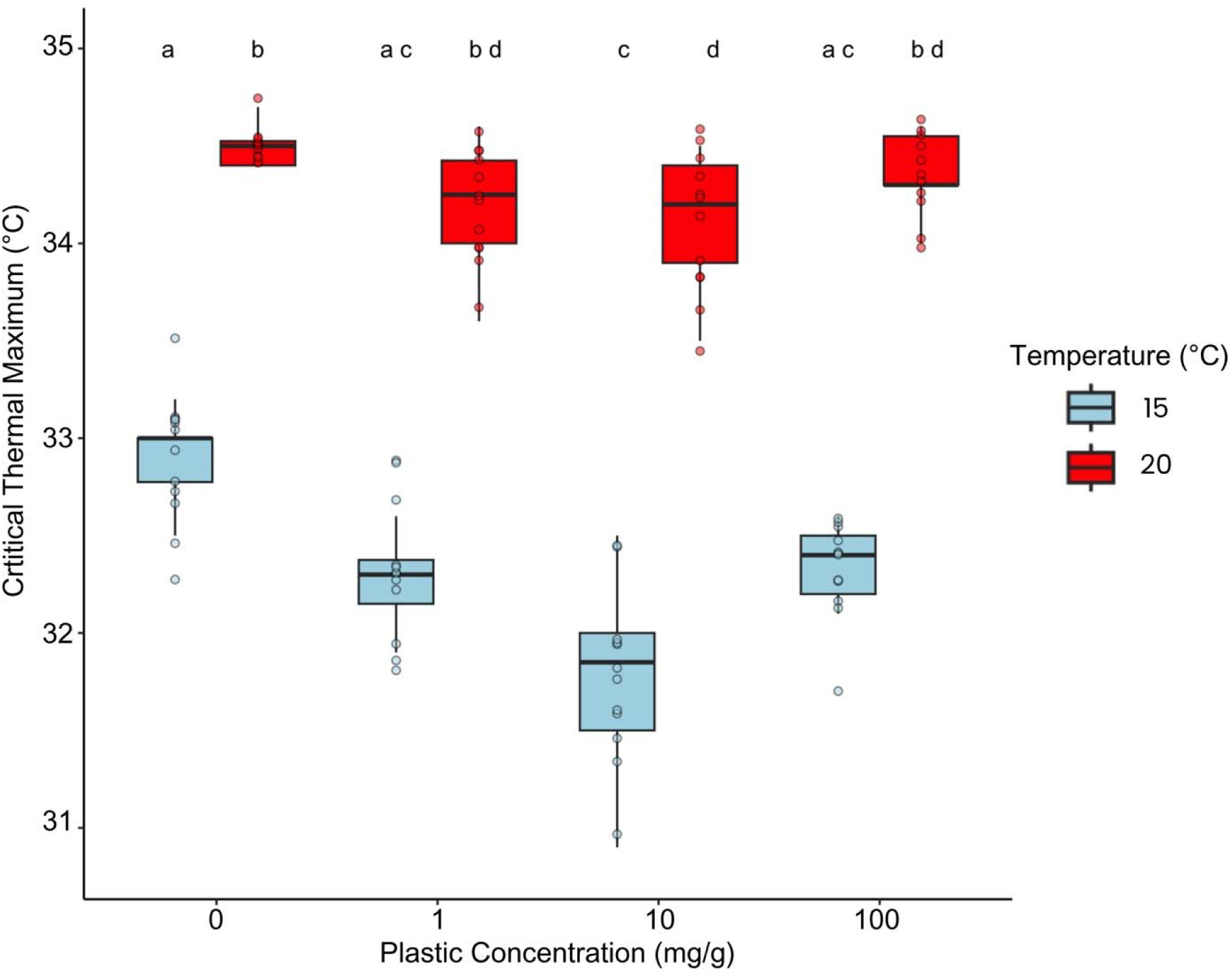
Boxplot of CTmax results for temperature (15°C and 20°C) and microplastic (0, 1, 10, 100 mg/g) treatments (n = 11-12), with individuals represented by dots. Bars represent 1.5x the interquartile range; letters indicate significance (Supplementary Tables 14-16).

**Table 2.**
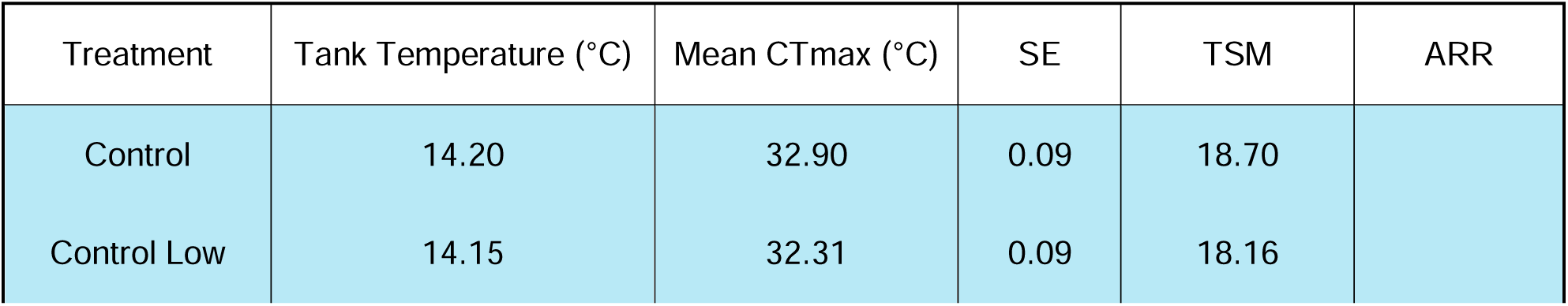
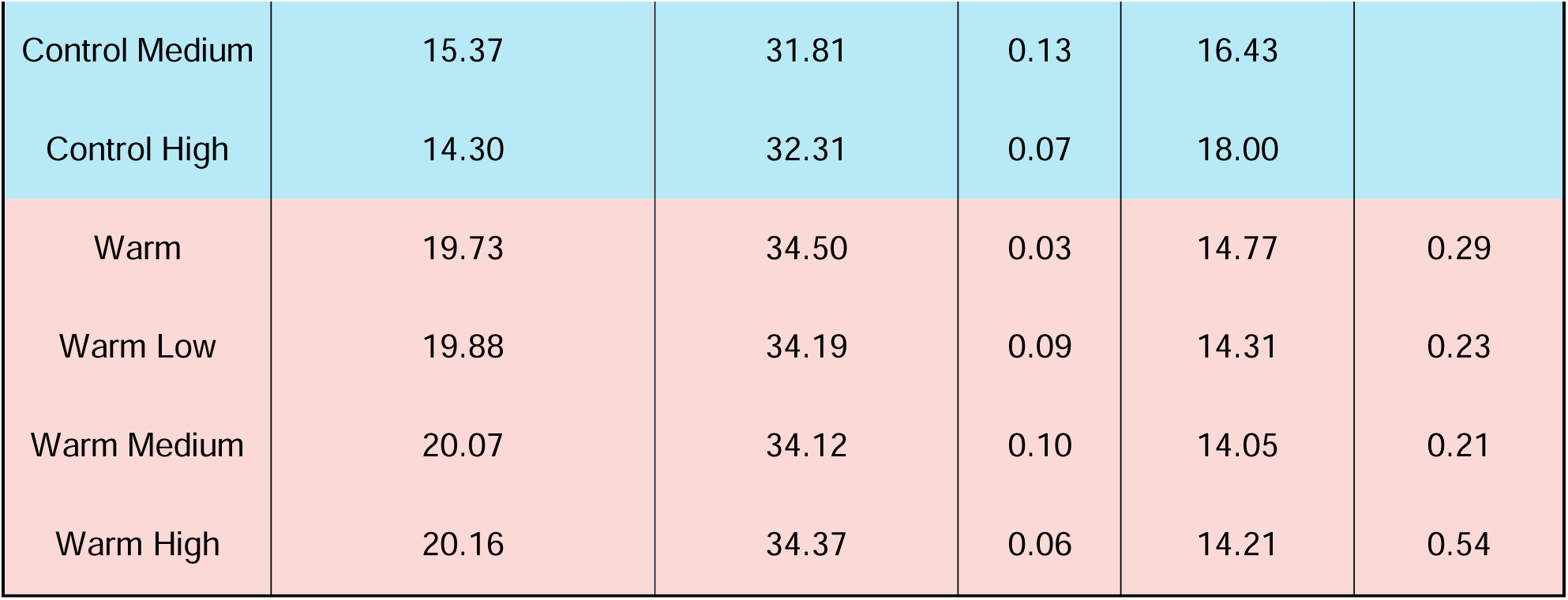
Summary of Critical Thermal Maximum values (mean + SE), Acclimation Response Ratios (ARR), and Thermal Safety Margins for temperature (15°C and 20°C) and microplastic (0, 1, 10, 100 mg/g) treatments (n = 11-12). See Supplementary Tables 14-16 for additional statistical information.

| Treatment | Tank Temperature (°C) | Mean CTmax (°C) | SE | TSM | ARR |
| --- | --- | --- | --- | --- | --- |
| Control | 14.20 | 32.90 | 0.09 | 18.70 |  |
| Control Low | 14.15 | 32.31 | 0.09 | 18.16 |  |
| Control Medium | 15.37 | 31.81 | 0.13 | 16.43 |  |
| Control High | 14.30 | 32.31 | 0.07 | 18.00 |  |
| Warm | 19.73 | 34.50 | 0.03 | 14.77 | 0.29 |
| Warm Low | 19.88 | 34.19 | 0.09 | 14.31 | 0.23 |
| Warm Medium | 20.07 | 34.12 | 0.10 | 14.05 | 0.21 |
| Warm High | 20.16 | 34.37 | 0.06 | 14.21 | 0.54 |

## DISCUSSION

Cumulative effects of temperature and plastic pollution are driving damaging and unprecedented effects on marine ecosystems and fishes. To characterize these effects, we exposed threespine stickleback to conditions of 15°C and 20°C (warming) and a gradient of nylon microplastic (0, 1, 10, and 100 mg nylon microplastic / g bloodworm) over 4 weeks, to investigate individual and combined impacts across behavioral, growth, and thermal physiological axes. Interestingly, we found plastics caused reductions in performance across these axes and while warming antagonistically interacted, leading to many associated trade-offs. Specifically, increasing plastic concentrations caused significant reductions in feeding rates, greater anxiety-associated behavior, and unique feeding behaviors, while warming temperatures had opposing effects. Plastic consumption likely drove many off-target effects (e.g., abnormal swelling of the gut and distended abdomens), particularly under warming conditions. Plastic exposure led to complex reductions in CT_max_ values, further reducing acclimation capacity, despite overall increases in thermal tolerance values (CT_max_) under warming conditions. Given these results, we anticipate greater anxiety behaviors and reductions in foraging efficiencies of fishes with increasing concentrations of plastic, which may be partially offset by warming. This offset however may drive greater lethal effects. Cumulatively, we expect these stressors yield greater energetic trade-offs, decreased accuracy in food selection, and reduce thermal acclimation capacities, with stark implications for marine ecosystem dynamics.

### Impacts of microplastics and warming during foraging impede energy acquisition

Feeding behavior defines energetic budgets and the key internalization pathway for external contaminants. Optimal foraging theory stipulates that animals act to maximize their energy acquisition during foraging (Pyke, 1984). The energetic costs and time spent searching for foraging habitat, handling, ingesting, and processing feed are weighed against their yield, with respect to an individual’s environmental and physiological constraints (Pyke, 1984). Variation in optimization within a generation or in response to changing conditions is dominantly dictated by changes in foraging behavior (e.g., changing foraging sites, learning, etc.) (Pyke, 1984; F. Zhang & Hui, 2014). We find impacts of warming and microplastics on the searching and consumption (handling and ingestion) stages of energy acquisition in fish.

A fish must explore its environment to effectively maximize feed while minimizing exposure to stress (e.g., predatory, temperature, pollutants). Previous work has demonstrated that warming temperatures contribute to exploration-exploitation trade-offs (Hajji & Lucas, 2026). This trade-off describes an individual’s decision framework to optimize resource gain and survival between exploration of known environments or exploitation of new environments without established risk (Berger-Tal et al., 2014; Mehlhorn et al., 2015). Inabilities to “choose” between environments leads to declines in resource acquisition and further survival. As warming temperature increased crosses to the white zone of the BWT assay, this work supports previous findings attributed to decreased problem solving and decisive commitments to crosses. Microplastics had opposing effects on crosses (i.e., were less likely to experience high cross counts), attributed to higher anxiety in the arena. Observed movements of fish away from the direct vicinity of food items following negative interactions with plastic particles (e.g., coughing) is indicative of an anxiety association. Such an association (i.e., microplastics as a negative reinforcer see Hineline, 1977) would contribute to rejections of foraging spaces, reducing time spent in foraging sites and feeding, while increasing search time. Inevitably, this would contribute to drastic declines foraging rates and efficiencies.

These stressors also have cumulative impacts on the consumption phases of feeding, which includes handling and ingestion. We found increasing microplastic concentrations yielded significant reductions in feeding rates (i.e., increased time foraging and consuming feed). Nylon microplastic have similarly reduced feeding rates in copepods on algae in a dose-dependent manner (Cole et al., 2019). With fibres representing the most abundant plastic form (70.6%) ingested by fish globally (Lim et al., 2022), these results have stark translations for fishes in natural ecosystems, with expected declines in feeding efficiencies. Intriguingly, when combined with warming, feeding rates partially recovered across plastic treatments, suggesting an antagonistic interaction. Antagonistic effects of warming on feeding rates may demand trade-offs and greater false selection of food items. Previous research has demonstrated that under warming conditions (exposure periods between 15-30 d), multiple fish species exhibit greater false selection of microplastic particles and carry higher microplastic body burdens (D’Avignon et al., 2023; Hasan et al., 2023; Wen et al., 2018). So, we suggest that through partially offsetting declines in feeding rates, warming drove greater false selection and increased ingestion of microplastic particles (i.e., increased microplastic body burdens; see *Abnormal morphologies and trade-offs in energetic yield with microplastic and warming)*. Further, after food is consumed, energy acquisition from feed may be additionally impeded due to microplastic contamination. Overall, reductions in feeding efficacy may have significant translations to energy consumption, predation risk, biomagnification, and selection of feeding sites (Gerking, 2014).

Alongside, microplastic contamination of feed yielded unique coughing and reselection behaviors that may have contributed to the observed decline in feeding rates. Other works have similarly observed coughing behaviors during microplastic feeding events (Li et al., 2021), which suggests some capacity for individual fish to reject ingestions after their initial false selection. This, however, still results in increased foraging durations and anxiety associated with feeding/prey selection (e.g., movement away from direct vicinity of food items, or towards corners of tank). We are unsure why fish reselected food particles “coughed up” by their conspecifics; we would suggest this may be due to social learning, imitation, and similar signalling that occurs when selecting a feeding site (Laland et al., 2011; Webster & Laland, 2018). Many fish species imitate diet selection and learn foraging behaviors (including foraging sites) from conspecifics (Brown & Laland, 2003; Pike et al., 2010). Particles rejected by conspecifics via coughing may be viewed as a safer alternative than exploring other feed available. We suspect the false selection of plastic particles over prey items may be due to their association and the false sense of satiety they provide individuals (Mahmood et al., 2024). Nonetheless, ingested microplastics are expected to contribute directly or indirectly (via energetic budgets) to declines in other performance factors.

Overall, we find that warming and microplastics likely are detrimental for and have compounding effects on energy acquisition. Under warming conditions, indecision between environments alongside greater microplastic ingestions, may drive significant reductions to energy acquisition. Inevitably, this would pose greater constraints to energetic budgets potentially hindering population fitness. As such, both plastic and temperature are considerable factors that should be applied within the optimal foraging theory framework, as reduced viability in prey and sites selection would decrease feeding performance, body condition, stress tolerance (e.g., thermal), and overall fitness.

### Abnormal morphologies and trade-offs in energetic yield with microplastic and warming

Growth metrics (body length, weight, condition factor) are key indicators of fitness representing energy assimilation and conversion from feeding after balancing allostatic load. Abnormalities are more common in polluted and stressed environments that disrupt genetic and physiological axes (Chandra et al., 2024; Gercken et al., 2006). While we did not observe significant differences in the weights nor lengths of fish, we observed a non-significant trend with plastic treatments demonstrating higher weights despite similar lengths. We attribute this potentially to increased retention of microplastic particles observed in other studies (Z. Wang et al., 2024), though larger sample sizes or longer exposure durations may be necessary to tease apart these differences. This is further supported by our own findings as fish under high plastic concentrations, especially with warming, exhibited apparent distention of the gut and abnormal swelling around the rectum. We suggest these effects may be due to 1) physical obstruction and distension of the gut by the plastic itself, and 2) gut dysbiosis. In regard to the former, blockages and distension change the relative position of their centre of mass and centre of buoyancy, driving instability and increased energetic costs (Webb & Weihs, 2015). The loss of dorsoventral positioning symptom was only observed under high plastic concentrations and warming and did not present under control temperatures. As such, under plastic exposure at higher temperatures fish may lack the energetic costs necessary to mitigate these symptoms due to other paramount energetic demands from allostatic load such as maintaining higher heat tolerance, as observed in the CT_max_ results (see discussion below). Other works have found similar results, with plastics causing blockages, lesions, dysbiosis, and swelling in the gut, which significantly reduces the fitness of an individual (Ali et al., 2023; Jin et al., 2018; Jovanović et al., 2018). Inability to maintain dorsoventral positioning would inevitably lead to lethality in the natural environment due to the inability to capture prey and evade predation, similarly to CT_max_ (Schulte, 2015).

### Plastics impede thermal performance

Ambient temperature is notably the dominate factor impacting thermal tolerance, however other stressors may impede thermal responses through various mechanisms (oxygen deficiencies, allostatic load and energetic costs, protein damage). Under current warming conditions, it has previously been established that marine stickleback populations are reaching their acclimation capacity (Hajji & Lucas, 2026). Our results support these findings, as warming temperatures, in isolation, yielded a lower TSM with an ARR value of 0.29 (compared reported values of 0.212 and 0.28), similarly indicating a compression of CT_max_ and unproportional shift in thermal tolerance (De Bonville et al., 2025; Hajji & Lucas, 2026). Though sublethal, greater energetic expenditure would be required under current warming conditions to reach similar performances as stickleback are outside their optimal thermal range and in the declines of their performance curve. This singular framework, however, did not consider how other stressors might cumulatively impede thermal performances and further contribute to energetic trade-offs, despite contemporary environmental conditions.

We found the addition of other microplastics complexly interacts to reduce thermal tolerance and acclimation capacity. Plastic exposure significantly decreased thermal tolerance values, with the 10 mg/g exposures having the lowest values for their respective temperature treatments, suggesting a biphasic thermal response. Surprisingly, the effects of plastic on CT_max_ were reduced under warming conditions. This may be due to a chronic upregulation of stress and heat factors to maintain functionality at higher temperatures (Metz et al., 2025), whereas under control temperature conditions energetic expenditure may prioritize other costs associated with the plastic exposure. Further, reductions in CT_max_ with plastic exposures was accompanied by lower TSM and ARR ratios, with a similar biphasic response was observed here. The seemingly biphasic thermal response within these concentrations of plastic exposures was unexpected, though we postulate many factors contribute.

The mechanism driving these effects is likely a combination of stress and energetic constraints. Previous works with copper exposure and cortisol ingestions have demonstrated similar reductions in CT_max_ (Bard et al., 2021; Crémazy et al., 2026), suggesting stress may play a role in this either through overlapping cortisol and heat shock factor mediated pathways or due to energetic constraints. The metabolic meltdown model may also play a role in this decrease, as previous studies have found reduced quality in feed drives declines in thermal performance (Huey & Kingsolver, 2019). Microplastic exposure may decrease energetic acquisition while increasing energy expenditure to combat these stressors, thereby increasing allostatic load and decreasing energy available for the acute thermal stress. It is unclear why a “rebound” effect, opposite of a hormetic response, was observed at the highest plastic concentration exposure across both temperatures (100 mg / g). Some research has demonstrated increases in energy acquisition capacity with greater declines in feed quality (Jobling, 1987). It may be possible that the highest concentrations drove such effects, increasing energy available during the acute thermal stress. The authors are unaware of any studies that have investigated these effects with microplastics and while beyond the scope of this study, we recommend future work should investigate how plastic exposure over shorter and longer durations of time and across difference acclimation temperatures, impacts the thermal performance of fish. Nonetheless, our results suggest areas with greater plastic pollution face additional thermal challenges and resilience constraints.

## CONCLUSION

As temperatures and plastic pollution levels in the ocean continue to rise, performance and function of marine animals are on declines. This work contributes to a growing body of literature emphasizing the importance of understanding cumulative effects by exposing fish to temperature conditions of 15°C and 20°C and microfibre plastic pollution across a concentration gradient (0-100 mg plastic/g food). We found plastics and warming interact complexly across different biological axes of fish. While warming offset some anxiety and reduced feeding efficiencies driven by microplastics, effects of plastic beyond a certain concentration threshold will continue to impede fish behavior. Despite antagonistic effects of warming, effects across other axes arise such as the morphological abnormalities (e.g., gut distension), which were prominently and lethally observed under cumulative stress. Of critical concern with increasing extreme climate events and warming conditions, fish are reaching the ends of their acclimatory capacity with plastics posing confounding thermal constraints. This raises critical concern for key fishery zones and may impede maximum sustainable yields, as heavily polluted areas (e.g., fishing grounds using nylon nets) and warming temperatures coincide spatially and temporarily with majority of annual commercial landings and the reproductive demographics of most fish populations. It is important to continue investigation of these cumulative effects and include additional factors (e.g., additives) to better predict long-term risks and compounding effects impeding ecosystem health. Understanding how these factors individually and in combination interact to disrupt fish health and performance, is of utmost importance to preventing marine ecosystem cascades, and directing management towards key vulnerable areas.

## ACKNOWLEDGEMENTS

Thank you to Sean Rogers, John Post, Gary Hardiman, and members of the Lucas, Rogers, and Jamniczky labs for helpful discussions. Thanks to BMSC and staff for support, and to Luke Anderson, Tao Eastham, Alessandra Gentile, and Amy Yuschychyn for their help with animal care. Thank you to Emily Atallah and Jian Jun Li of the UCalgary Faculty of Science Chemistry Department and Instrumentation Facility for their help in conducting FTIR analysis. This work was supported by a Natural Sciences and Research Council of Canada (NSERC) Discovery Grant and Discovery Launch Supplement, a Canada Foundation for Innovation John R. Evans Leaders Fund award, and UCalgary VPR Catalyst Award and start-up funding to KNL. ALH was also supported by an Alberta Graduate Excellence Scholarship, Maritime Awards Society of Canada Graduate Scholarship, and Alumni Association Graduate Scholarship through UCalgary. The authors declare no competing or financial interests.

## SUPPLEMENTARY

**Supplementary Table 1.**
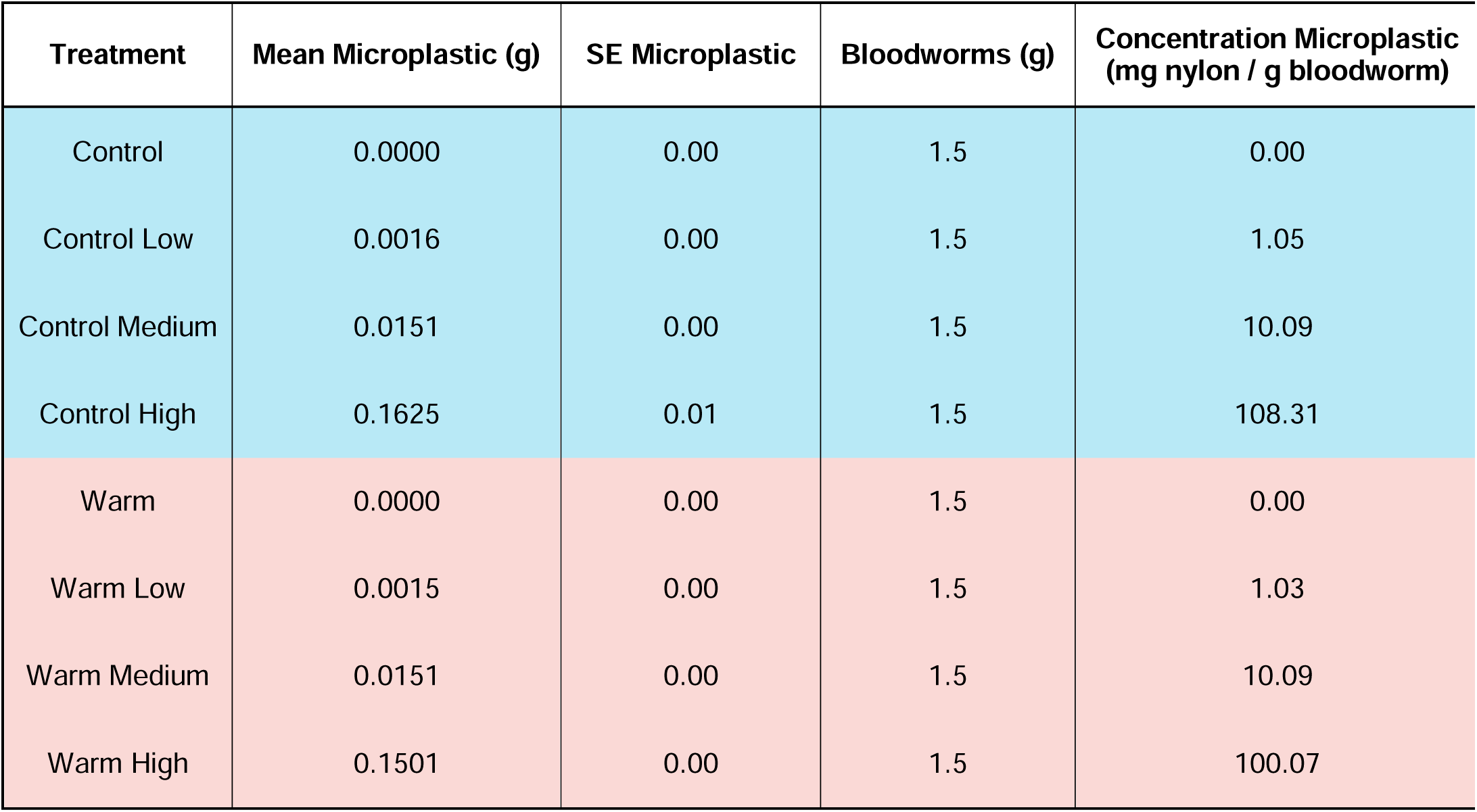
Tank feed data for temperature (15°C and 20°C) and microplastic (0, 1, 10, 100 mg/g) treatments.

**Supplementary Table 2.**
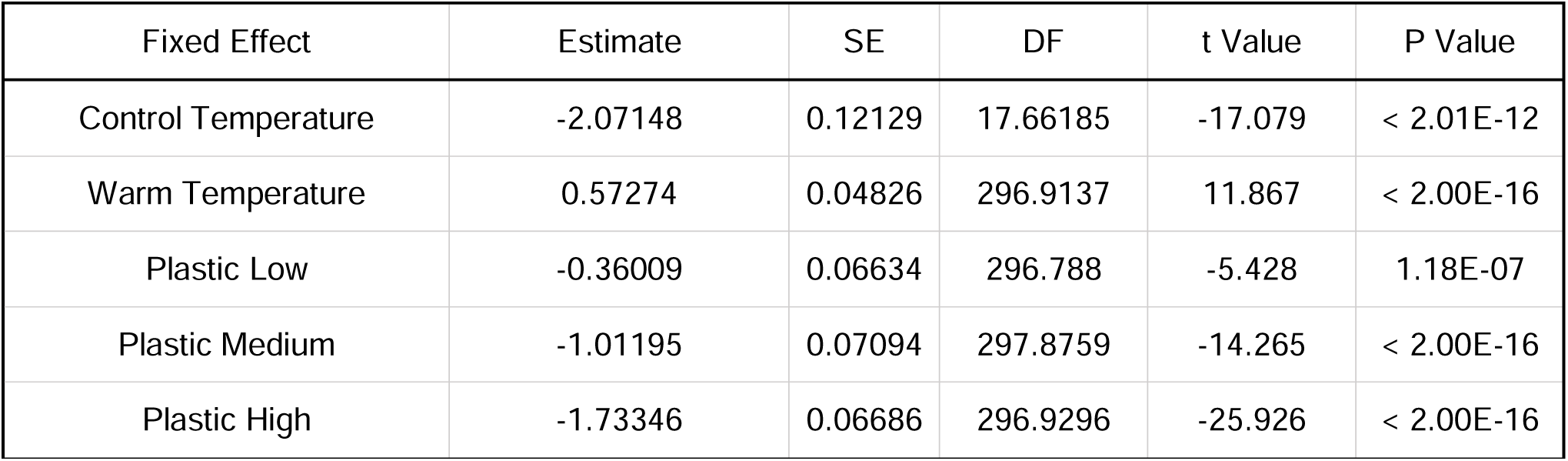
LMER statistical summary of tank feeding rates (log(mg/s)) for temperature (15°C and 20°C) and microplastic (0, 1, 10, 100 mg/g) treatments (n = 32-42).

**Supplementary Table 3.**
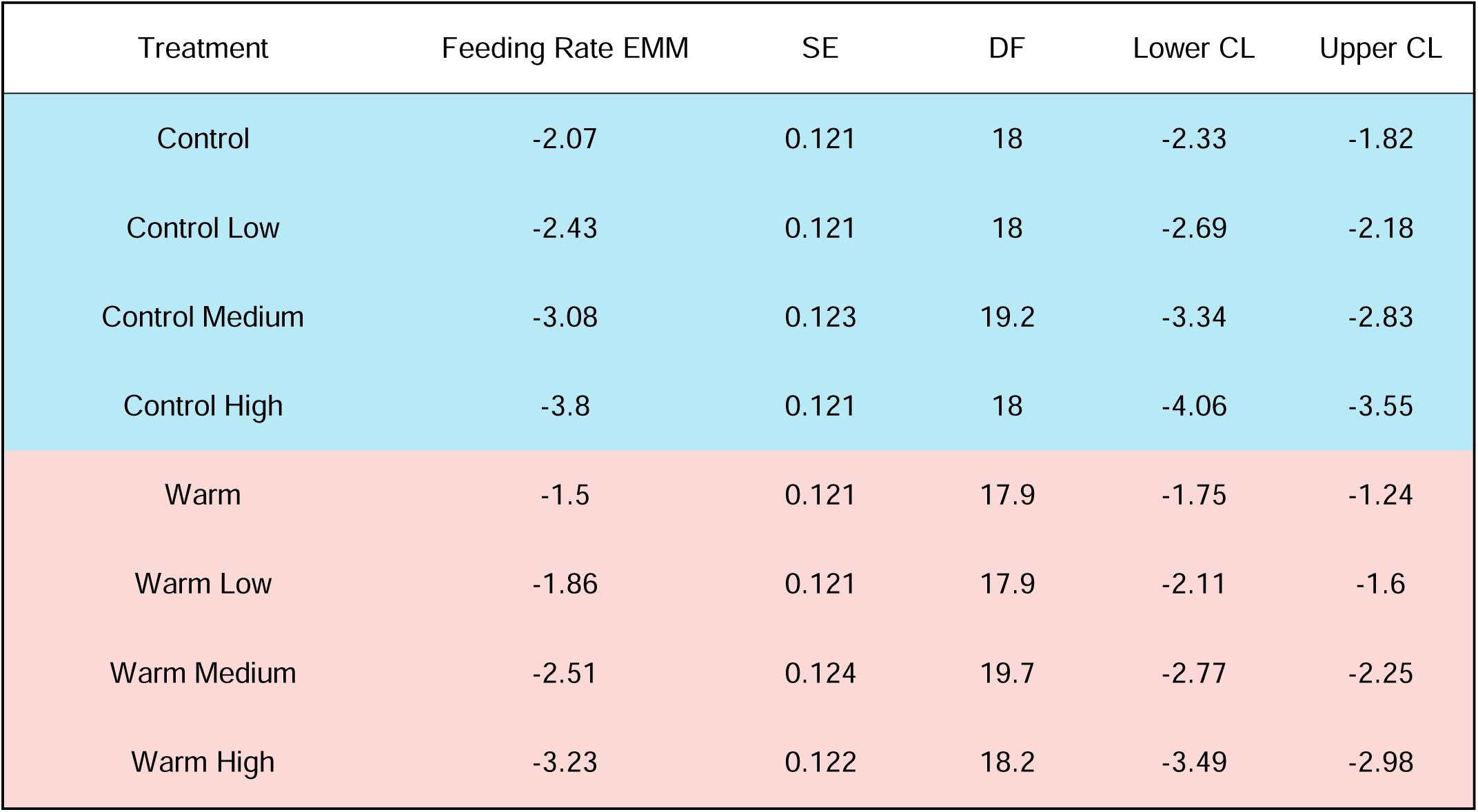
Estimated marginal mean tank feeding rates (log(mg/s)) summary table for temperature (15°C and 20°C) and microplastic (0, 1, 10, 100 mg/g) treatments (n = 32-42).

**Supplementary Table 4.**
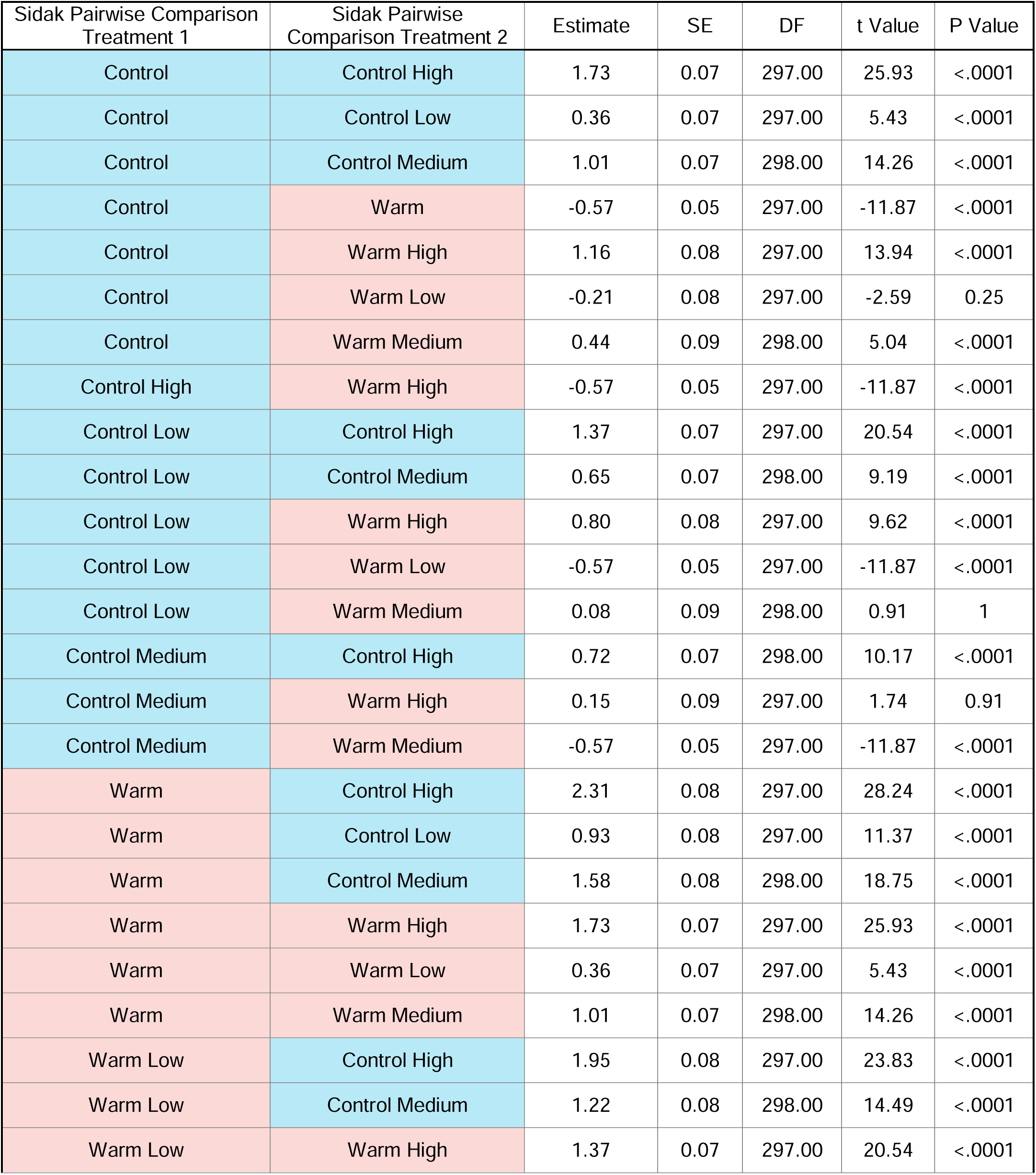
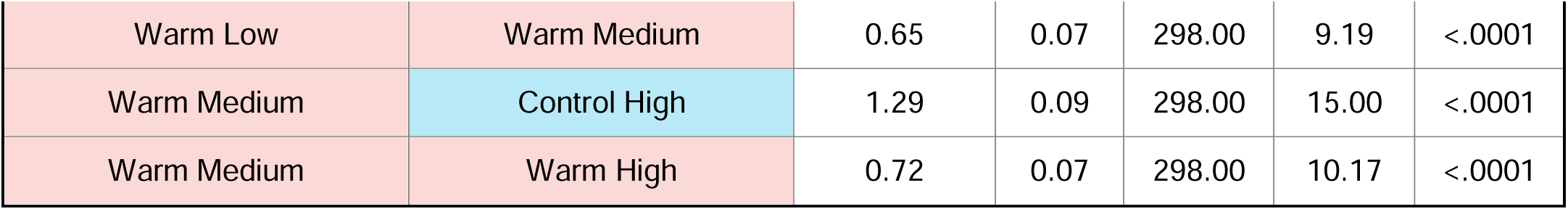
Sidak comparisons of tank feeding rates (log(mg/s)) for temperature (15°C and 20°C) and microplastic (0, 1, 10, 100 mg/g) treatments (n = 32-42).

**Supplementary Table 5.**
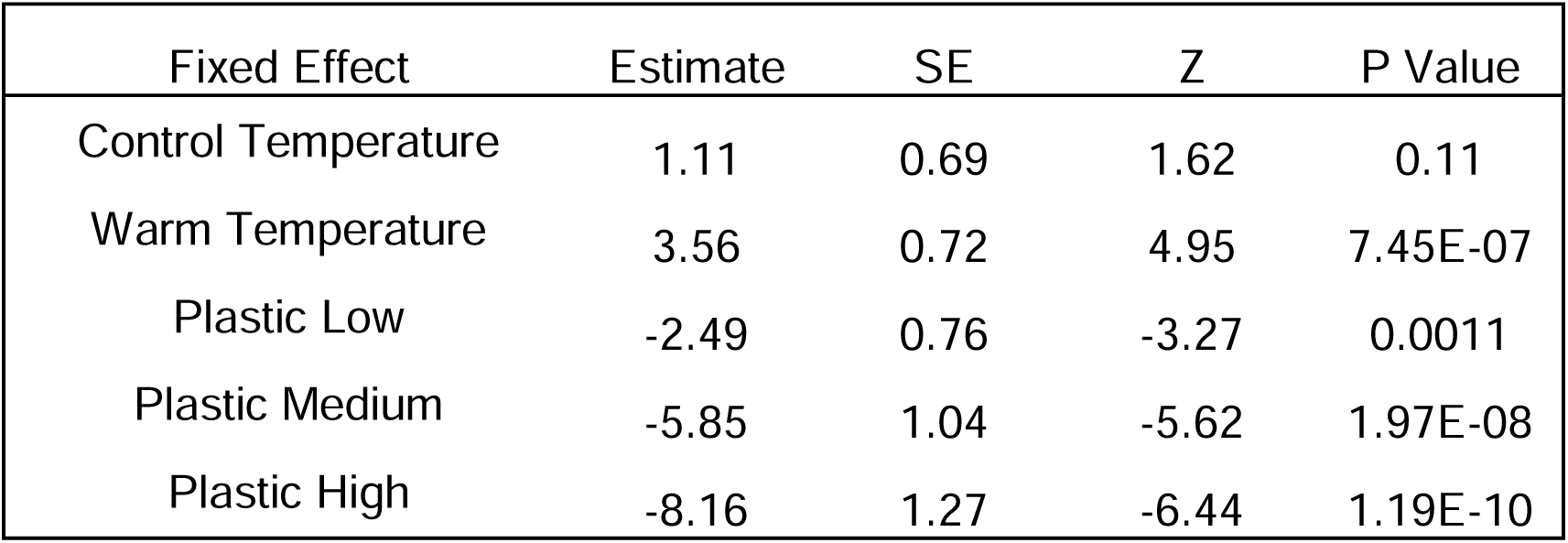
GLMER statistical summary for feeding rates used in the OR calculations with tank and day as nested effects for temperature (15°C and 20°C) and microplastic (0, 1, 10, 100 mg/g) treatments (n = 32-42).

**Supplementary Table 6.**
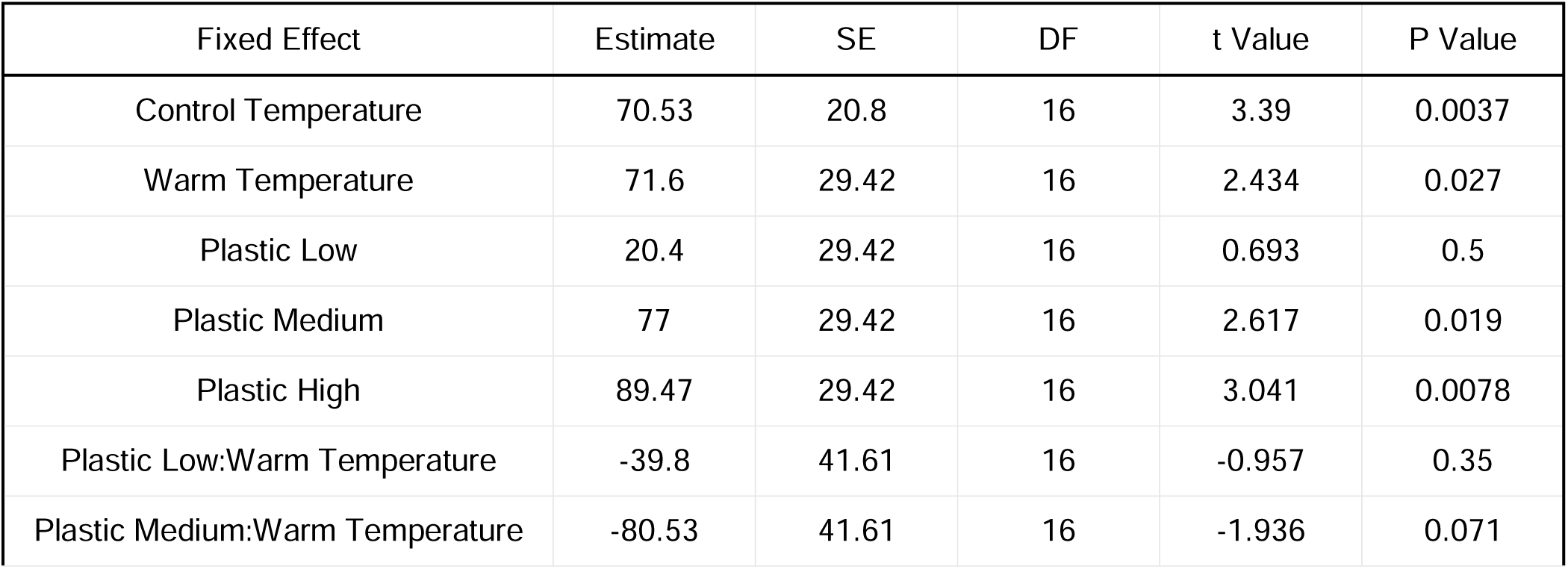
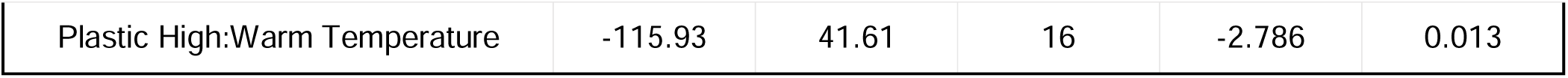
LMER statistical summary for Black White Test duration spent in white zone for temperature (15°C and 20°C) and microplastic (0, 1, 10, 100 mg/g) treatments (n = 15).

**Supplementary Table 7.**
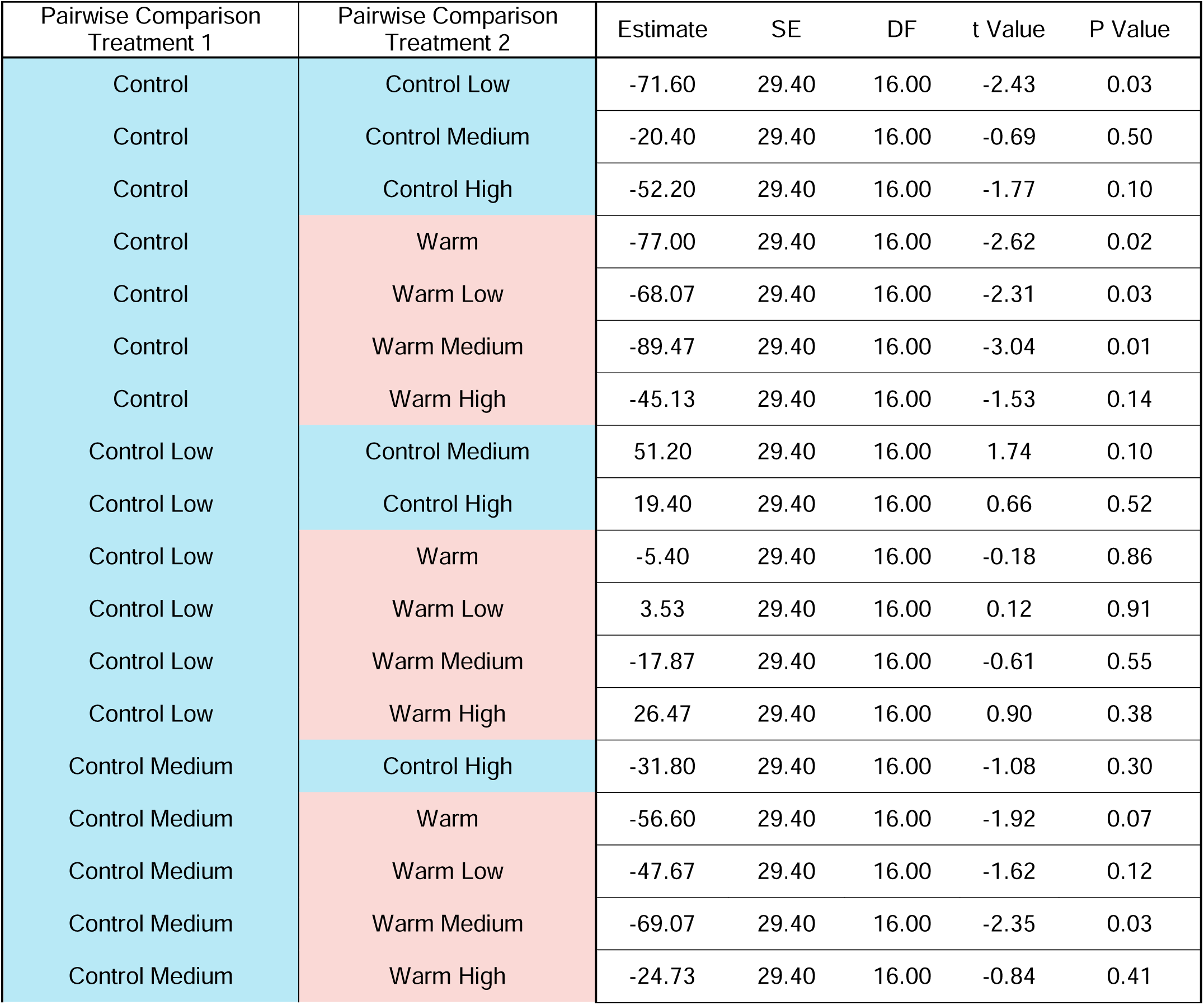
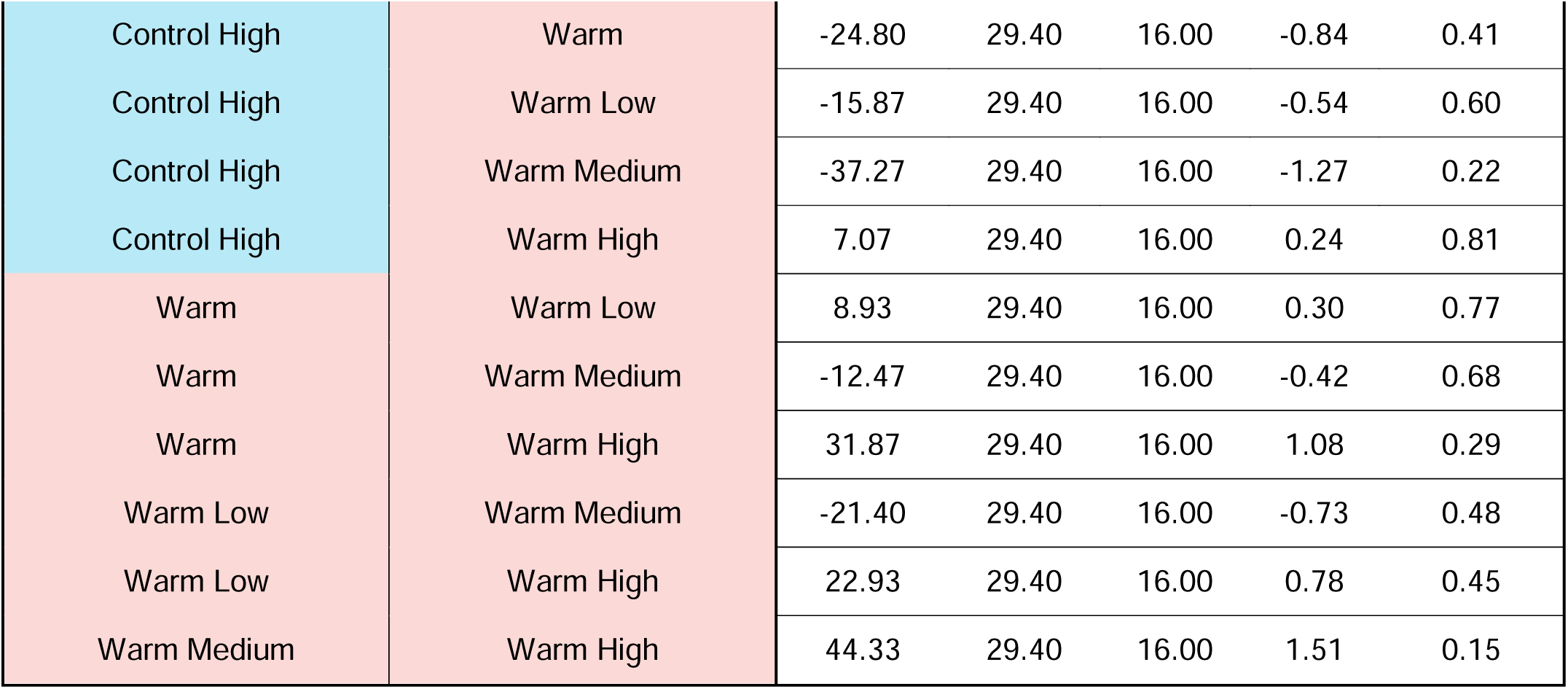
Pairwise comparisons for Black White Test duration spent in white zone for temperature (15°C and 20°C) and microplastic (0, 1, 10, 100 mg/g) treatments (n = 15).

**Supplementary Table 8.**
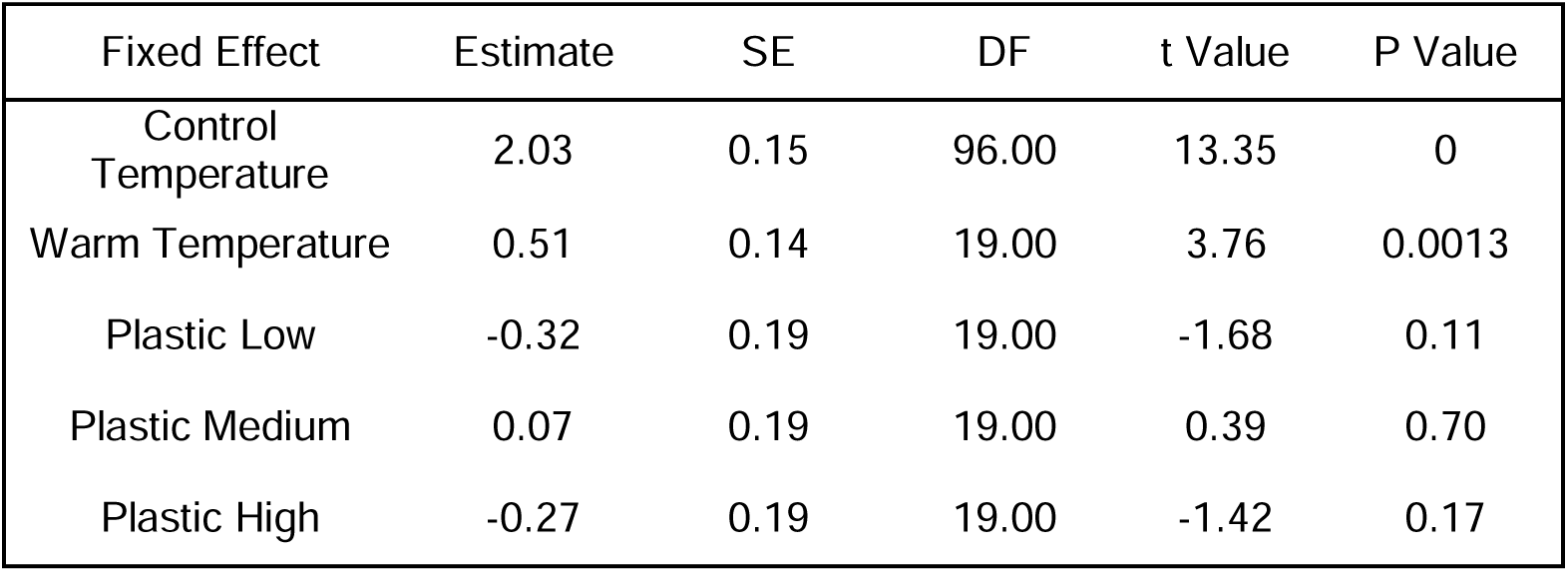
GAM statistical summary for Black White Test crosses to the white zone for temperature (15°C and 20°C) and microplastic (0, 1, 10, 100 mg/g) treatments (n = 15).

**Supplementary Table 9.**
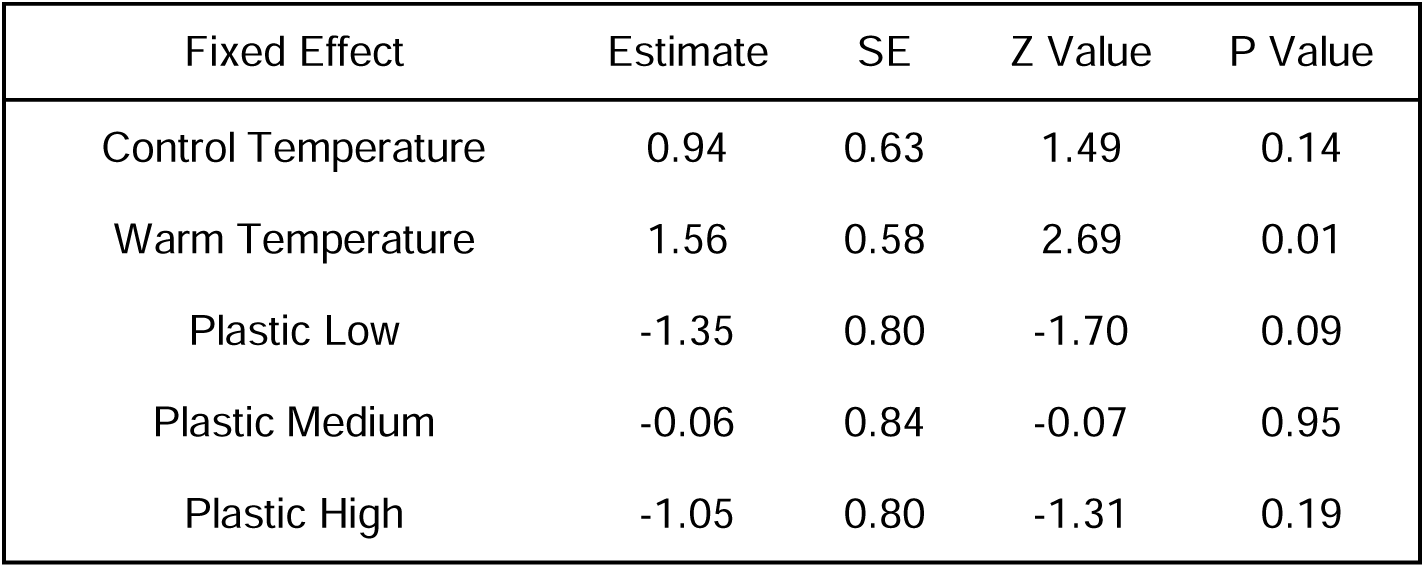
GLMER statistical summary for Black White Test frequency of visits to the white zone used in the OR calculations for temperature (15°C and 20°C) and microplastic (0, 1, 10, 100 mg/g) treatments (n = 15).

**Supplementary Table 10.**
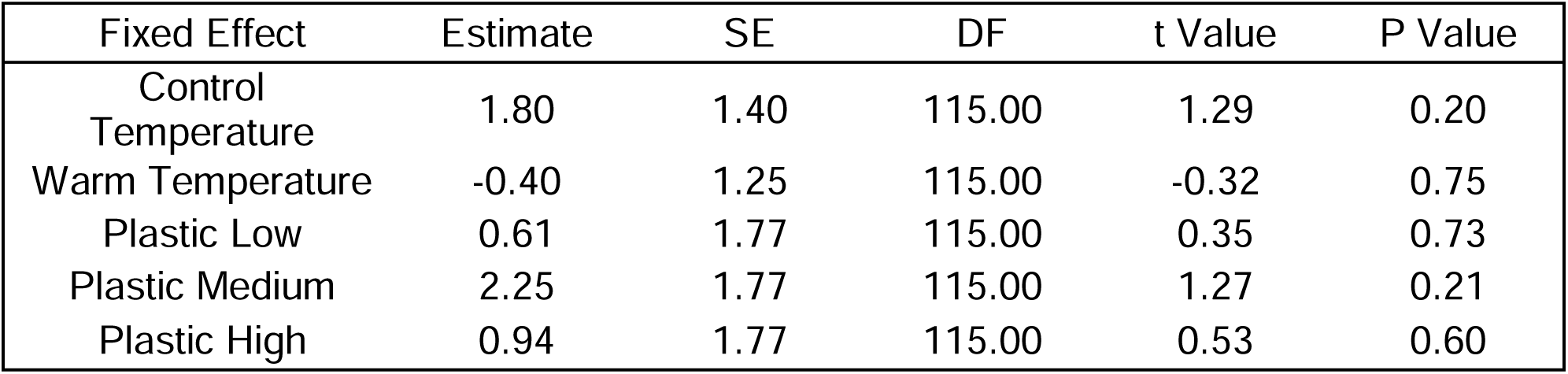
LMER statistical summary for individual change in length for temperature (15°C and 20°C) and microplastic (0, 1, 10, 100 mg/g) treatments (n = 15).

**Supplementary Table 11.**
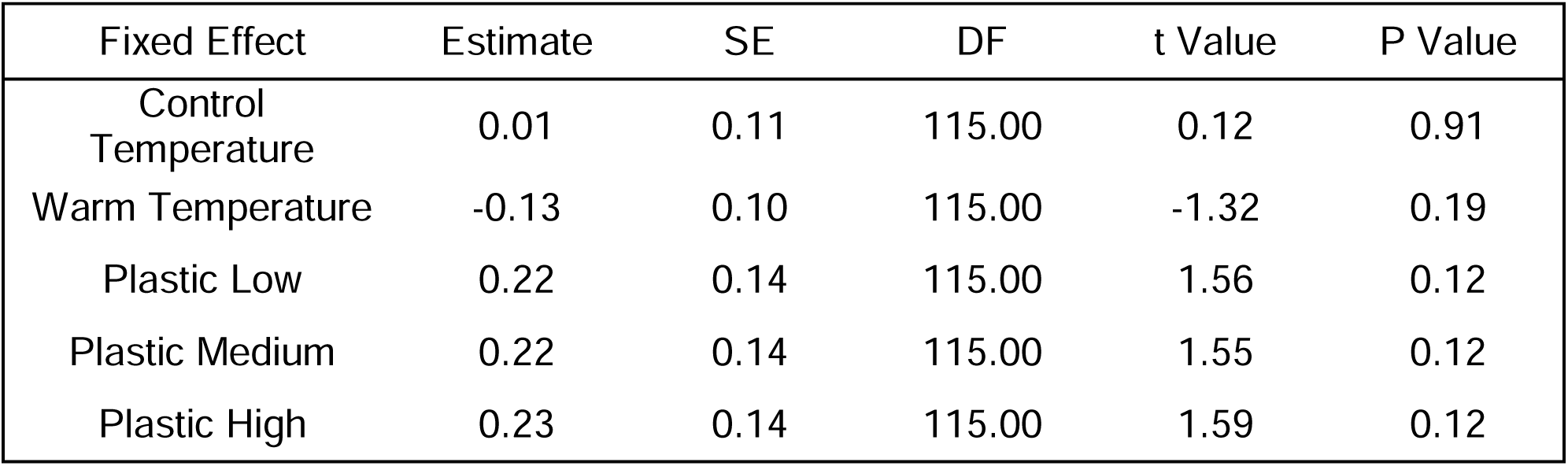
LMER statistical summary for individual change in weight for temperature (15°C and 20°C) and microplastic (0, 1, 10, 100 mg/g) treatments (n = 15).

**Supplementary Table 12.**
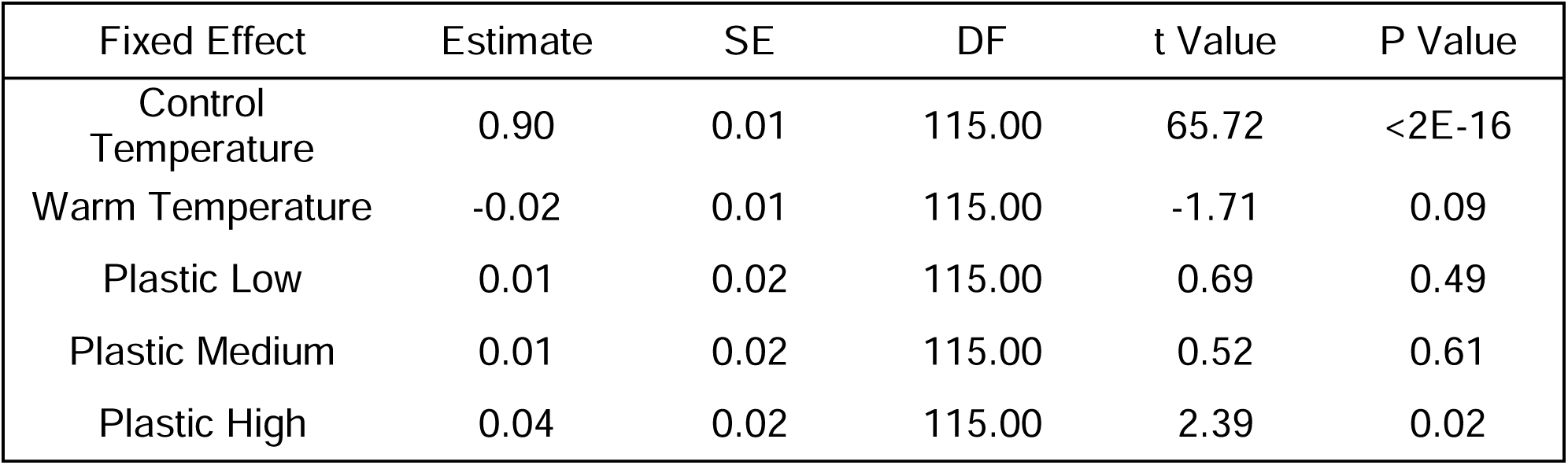
LMER statistical summary for Condition Factor for temperature (15°C and 20°C) and microplastic (0, 1, 10, 100 mg/g) treatments (n = 15).

**Supplementary Table 13.**
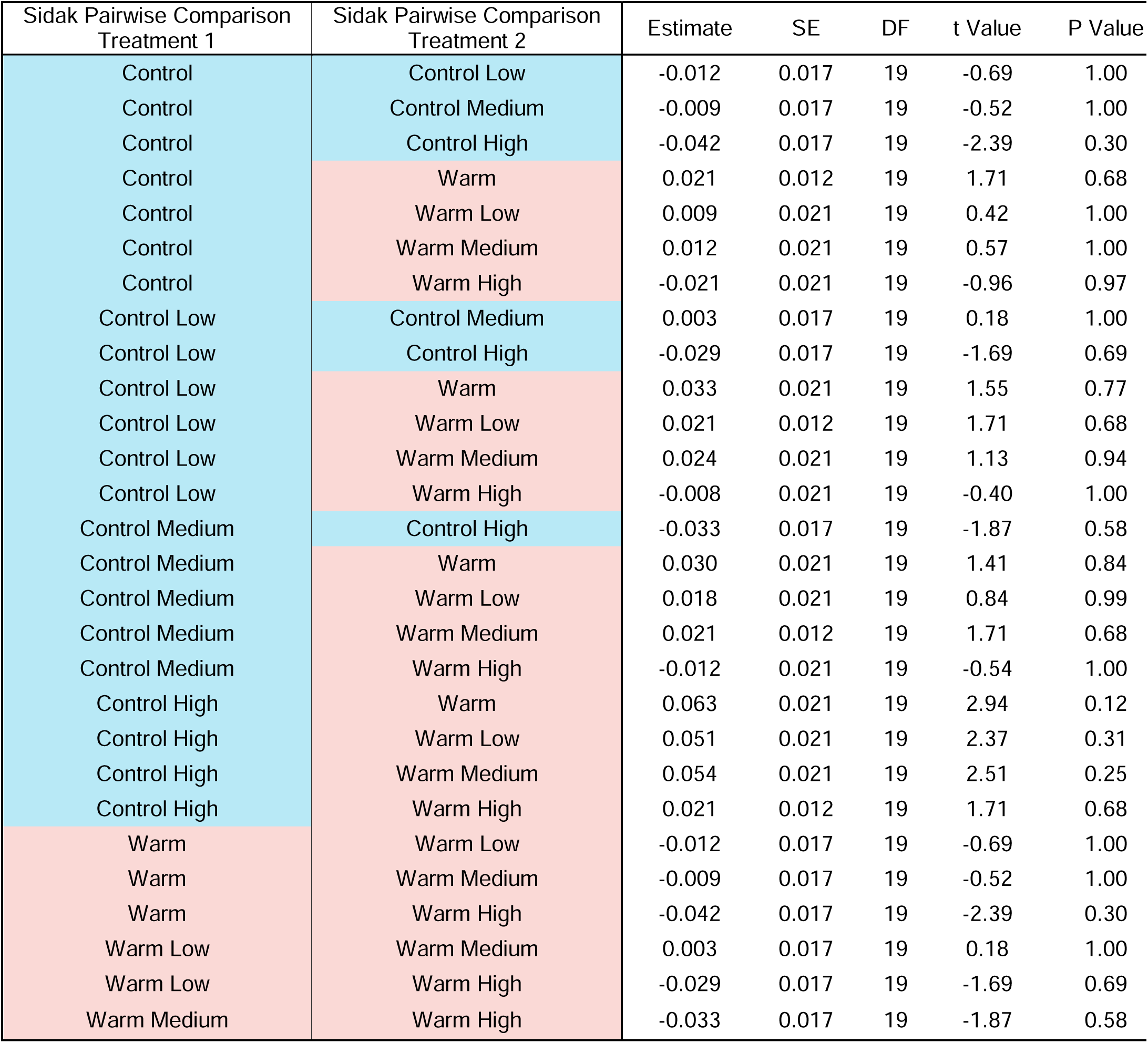
Sidak comparisons of condition factor for temperature (15°C and 20°C) and microplastic (0, 1, 10, 100 mg/g) treatments (n = 15).

**Supplementary Table 14.**
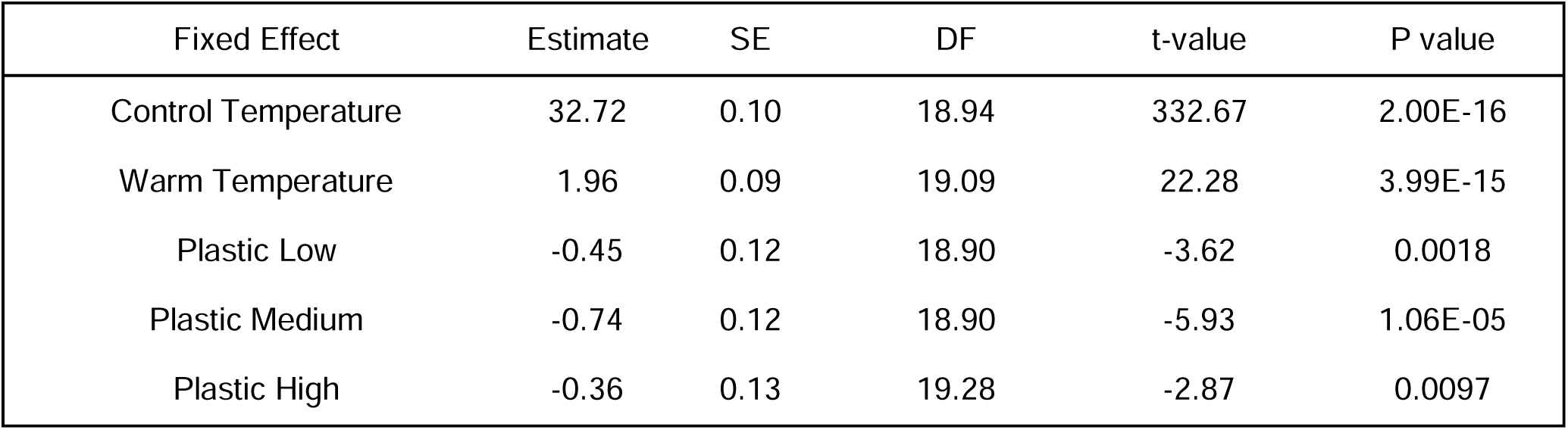
LMER statistical summary of Critical Thermal Maximum for temperature (15°C and 20°C) and microplastic (0, 1, 10, 100 mg/g) treatments (n = 11-12). Temperature and Plastic as independent fixed effects and tanks as nested factor.

**Supplementary Table 15.**
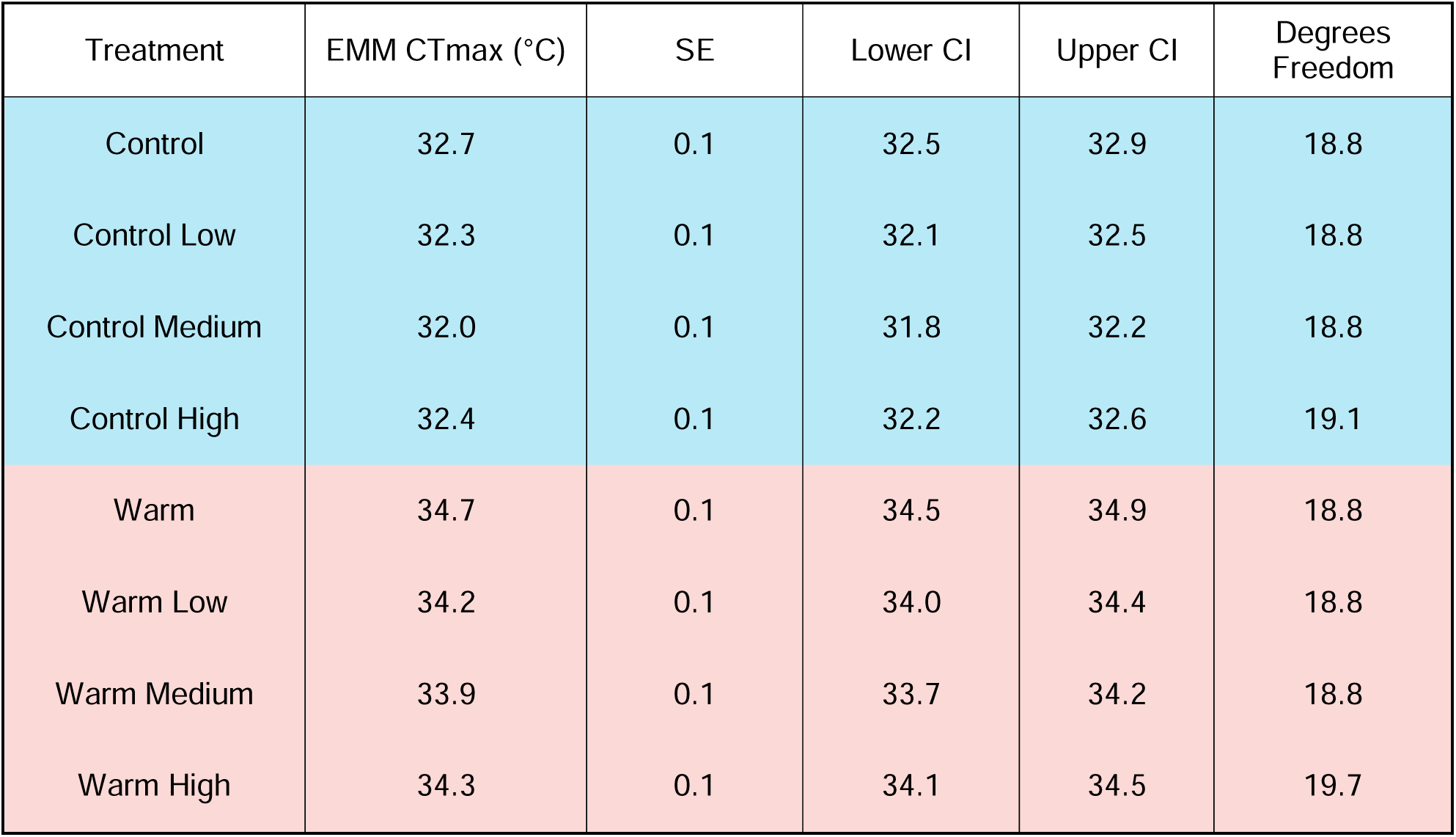
Estimated marginal mean tank Critical Thermal Maximum summary table for temperature (15°C and 20°C) and microplastic (0, 1, 10, 100 mg/g) treatments (n = 11- 12).

**Supplementary Table 16.**
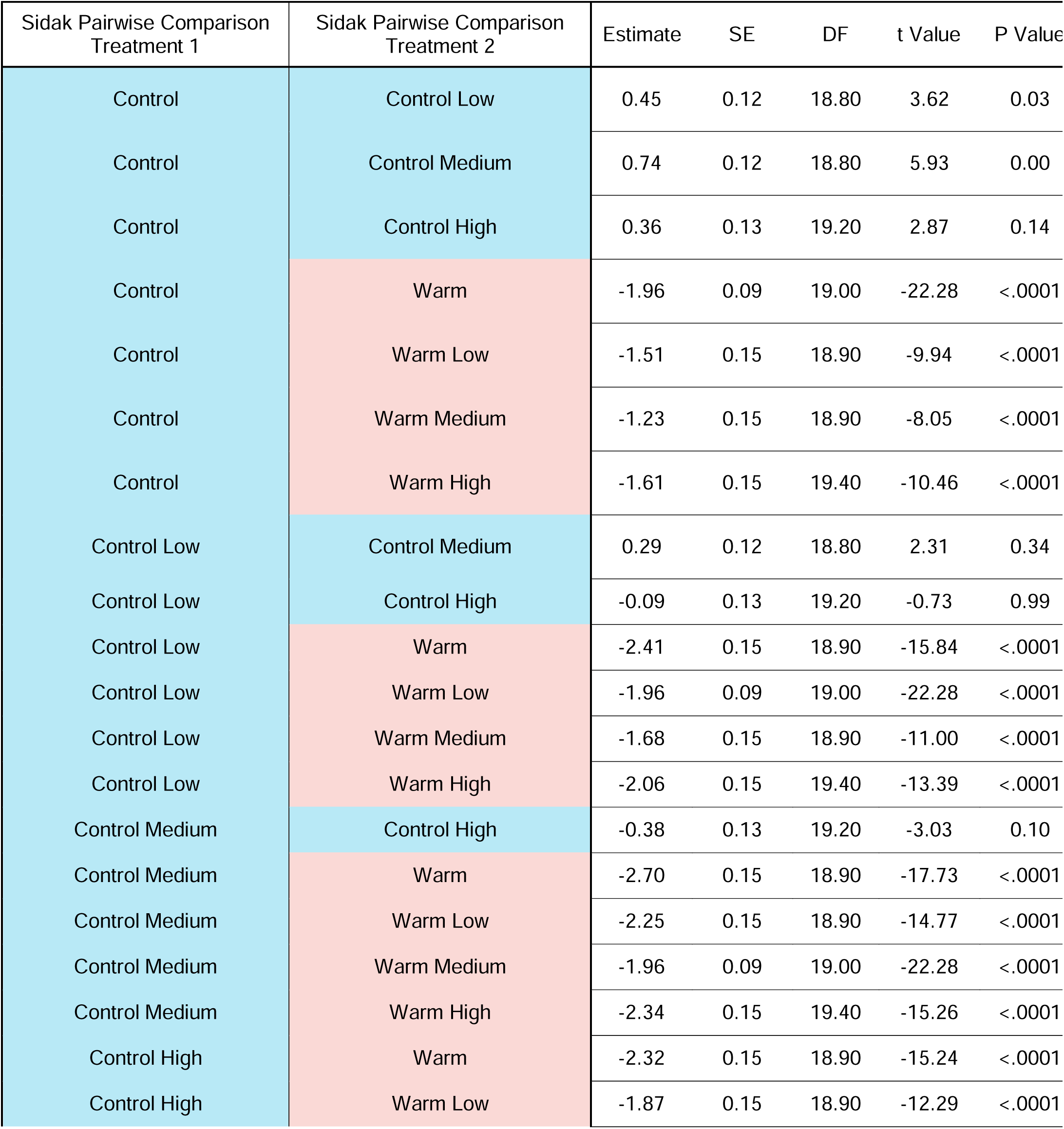
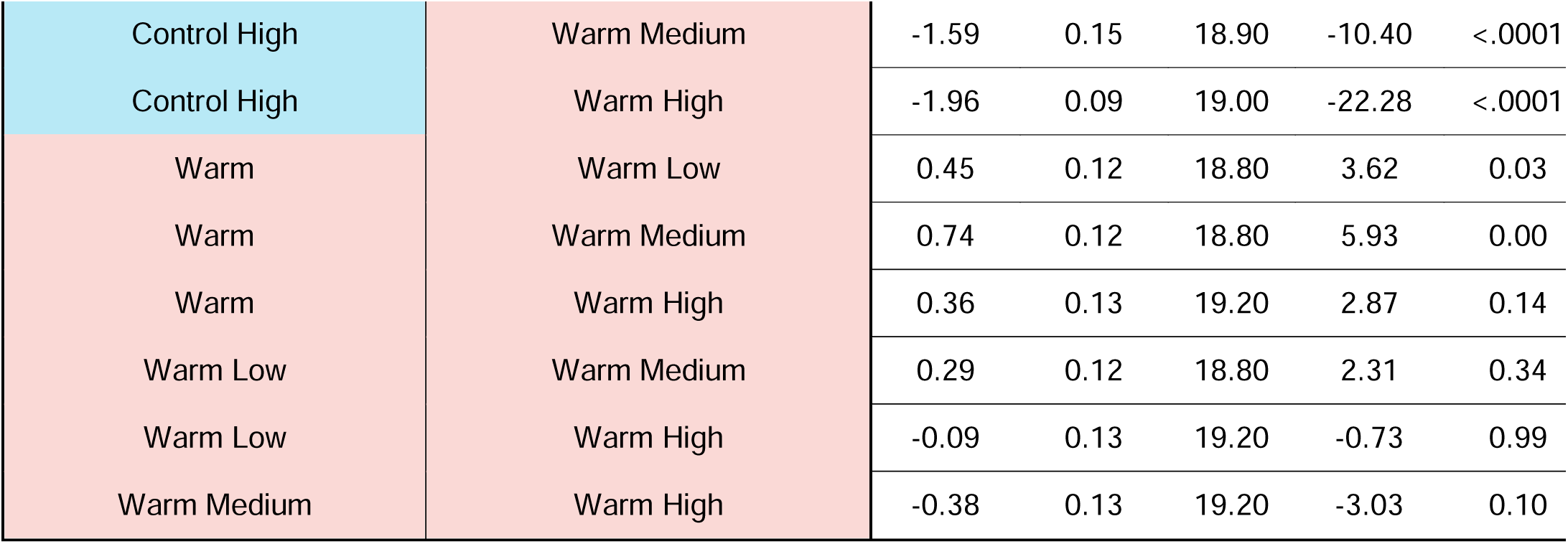
Sidak comparisons of Critical Thermal Maximum values for temperature (15°C and 20°C) and microplastic (0, 1, 10, 100 mg/g) treatments (n = 11-12).

