## Supplementary Material for "Cumulative effects of nylon microplastic fibres and warming temperature on the behavior and physiology of marine threespine stickleback (*Gasterosteus aculeatus*)"

Supplementary Table 1. Tank feed data for temperature (15°C and 20°C) and microplastic (0, 1, 10, 100 mg/g) treatments.

| **Treatment** | **Mean Microplastic (g)** | **SE Microplastic** | **Bloodworms (g)** | **Concentration Microplastic  (mg nylon / g bloodworm)** |
| --- | --- | --- | --- | --- |
| Control | 0.0000 | 0.00 | 1.5 | 0.00 |
| Control Low | 0.0016 | 0.00 | 1.5 | 1.05 |
| Control Medium | 0.0151 | 0.00 | 1.5 | 10.09 |
| Control High | 0.1625 | 0.01 | 1.5 | 108.31 |
| Warm | 0.0000 | 0.00 | 1.5 | 0.00 |
| Warm Low | 0.0015 | 0.00 | 1.5 | 1.03 |
| Warm Medium | 0.0151 | 0.00 | 1.5 | 10.09 |
| Warm High | 0.1501 | 0.00 | 1.5 | 100.07 |

Supplementary Table 2. LMER statistical summary of tank feeding rates (log(mg/s)) for temperature (15°C and 20°C) and microplastic (0, 1, 10, 100 mg/g) treatments (n = 32-42).

| Fixed Effect | Estimate | SE | DF | t Value | P Value |
| --- | --- | --- | --- | --- | --- |
| Control Temperature | -2.07148 | 0.12129 | 17.66185 | -17.079 | < 2.01E-12 |
| Warm Temperature | 0.57274 | 0.04826 | 296.9137 | 11.867 | < 2.00E-16 |
| Plastic Low | -0.36009 | 0.06634 | 296.788 | -5.428 | 1.18E-07 |
| Plastic Medium | -1.01195 | 0.07094 | 297.8759 | -14.265 | < 2.00E-16 |
| Plastic High | -1.73346 | 0.06686 | 296.9296 | -25.926 | < 2.00E-16 |

Supplementary Table 3. Estimated marginal mean tank feeding rates (log(mg/s)) summary table for temperature (15°C and 20°C) and microplastic (0, 1, 10, 100 mg/g) treatments (n = 32-42).

| Treatment | Feeding Rate EMM | SE | DF | Lower CL | Upper CL |
| --- | --- | --- | --- | --- | --- |
| Control | -2.07 | 0.121 | 18 | -2.33 | -1.82 |
| Control Low | -2.43 | 0.121 | 18 | -2.69 | -2.18 |
| Control Medium | -3.08 | 0.123 | 19.2 | -3.34 | -2.83 |
| Control High | -3.8 | 0.121 | 18 | -4.06 | -3.55 |
| Warm | -1.5 | 0.121 | 17.9 | -1.75 | -1.24 |
| Warm Low | -1.86 | 0.121 | 17.9 | -2.11 | -1.6 |
| Warm Medium | -2.51 | 0.124 | 19.7 | -2.77 | -2.25 |
| Warm High | -3.23 | 0.122 | 18.2 | -3.49 | -2.98 |

Supplementary Table 4. Sidak comparisons of tank feeding rates (log(mg/s)) for temperature (15°C and 20°C) and microplastic (0, 1, 10, 100 mg/g) treatments (n = 32-42).

| Sidak Pairwise Comparison Treatment 1 | Sidak Pairwise Comparison Treatment 2 | Estimate | SE | DF | t Value | P Value |
| --- | --- | --- | --- | --- | --- | --- |
| Control | Control High | 1.73 | 0.07 | 297.00 | 25.93 | <.0001 |
| Control | Control Low | 0.36 | 0.07 | 297.00 | 5.43 | <.0001 |
| Control | Control Medium | 1.01 | 0.07 | 298.00 | 14.26 | <.0001 |
| Control | Warm | -0.57 | 0.05 | 297.00 | -11.87 | <.0001 |
| Control | Warm High | 1.16 | 0.08 | 297.00 | 13.94 | <.0001 |
| Control | Warm Low | -0.21 | 0.08 | 297.00 | -2.59 | 0.25 |
| Control | Warm Medium | 0.44 | 0.09 | 298.00 | 5.04 | <.0001 |
| Control High | Warm High | -0.57 | 0.05 | 297.00 | -11.87 | <.0001 |
| Control Low | Control High | 1.37 | 0.07 | 297.00 | 20.54 | <.0001 |
| Control Low | Control Medium | 0.65 | 0.07 | 298.00 | 9.19 | <.0001 |
| Control Low | Warm High | 0.80 | 0.08 | 297.00 | 9.62 | <.0001 |
| Control Low | Warm Low | -0.57 | 0.05 | 297.00 | -11.87 | <.0001 |
| Control Low | Warm Medium | 0.08 | 0.09 | 298.00 | 0.91 | 1 |
| Control Medium | Control High | 0.72 | 0.07 | 298.00 | 10.17 | <.0001 |
| Control Medium | Warm High | 0.15 | 0.09 | 297.00 | 1.74 | 0.91 |
| Control Medium | Warm Medium | -0.57 | 0.05 | 297.00 | -11.87 | <.0001 |
| Warm | Control High | 2.31 | 0.08 | 297.00 | 28.24 | <.0001 |
| Warm | Control Low | 0.93 | 0.08 | 297.00 | 11.37 | <.0001 |
| Warm | Control Medium | 1.58 | 0.08 | 298.00 | 18.75 | <.0001 |
| Warm | Warm High | 1.73 | 0.07 | 297.00 | 25.93 | <.0001 |
| Warm | Warm Low | 0.36 | 0.07 | 297.00 | 5.43 | <.0001 |
| Warm | Warm Medium | 1.01 | 0.07 | 298.00 | 14.26 | <.0001 |
| Warm Low | Control High | 1.95 | 0.08 | 297.00 | 23.83 | <.0001 |
| Warm Low | Control Medium | 1.22 | 0.08 | 298.00 | 14.49 | <.0001 |
| Warm Low | Warm High | 1.37 | 0.07 | 297.00 | 20.54 | <.0001 |
| Warm Low | Warm Medium | 0.65 | 0.07 | 298.00 | 9.19 | <.0001 |
| Warm Medium | Control High | 1.29 | 0.09 | 298.00 | 15.00 | <.0001 |
| Warm Medium | Warm High | 0.72 | 0.07 | 298.00 | 10.17 | <.0001 |

Supplementary Table 5. GLMER statistical summary for feeding rates used in the OR calculations with tank and day as nested effects for temperature (15°C and 20°C) and microplastic (0, 1, 10, 100 mg/g) treatments (n = 32-42).

| Fixed Effect | Estimate | SE | Z | P Value |
| --- | --- | --- | --- | --- |
| Control Temperature | 1.11 | 0.69 | 1.62 | 0.11 |
| Warm Temperature | 3.56 | 0.72 | 4.95 | 7.45E-07 |
| Plastic Low | -2.49 | 0.76 | -3.27 | 0.0011 |
| Plastic Medium | -5.85 | 1.04 | -5.62 | 1.97E-08 |
| Plastic High | -8.16 | 1.27 | -6.44 | 1.19E-10 |

Supplementary Table 6. LMER statistical summary for Black White Test duration spent in white zone for temperature (15°C and 20°C) and microplastic (0, 1, 10, 100 mg/g) treatments (n = 15).

| Fixed Effect | Estimate | SE | DF | t Value | P Value |
| --- | --- | --- | --- | --- | --- |
| Control Temperature | 70.53 | 20.8 | 16 | 3.39 | 0.0037 |
| Warm Temperature | 71.6 | 29.42 | 16 | 2.434 | 0.027 |
| Plastic Low | 20.4 | 29.42 | 16 | 0.693 | 0.5 |
| Plastic Medium | 77 | 29.42 | 16 | 2.617 | 0.019 |
| Plastic High | 89.47 | 29.42 | 16 | 3.041 | 0.0078 |
| Plastic Low:Warm Temperature | -39.8 | 41.61 | 16 | -0.957 | 0.35 |
| Plastic Medium:Warm Temperature | -80.53 | 41.61 | 16 | -1.936 | 0.071 |
| Plastic High:Warm Temperature | -115.93 | 41.61 | 16 | -2.786 | 0.013 |

Supplementary Table 7. Pairwise comparisons for Black White Test duration spent in white zone for temperature (15°C and 20°C) and microplastic (0, 1, 10, 100 mg/g) treatments (n = 15).

| Pairwise Comparison Treatment 1 | Pairwise Comparison Treatment 2 | Estimate | SE | DF | t Value | P Value |
| --- | --- | --- | --- | --- | --- | --- |
| Control | Control Low | -71.60 | 29.40 | 16.00 | -2.43 | 0.03 |
| Control | Control Medium | -20.40 | 29.40 | 16.00 | -0.69 | 0.50 |
| Control | Control High | -52.20 | 29.40 | 16.00 | -1.77 | 0.10 |
| Control | Warm | -77.00 | 29.40 | 16.00 | -2.62 | 0.02 |
| Control | Warm Low | -68.07 | 29.40 | 16.00 | -2.31 | 0.03 |
| Control | Warm Medium | -89.47 | 29.40 | 16.00 | -3.04 | 0.01 |
| Control | Warm High | -45.13 | 29.40 | 16.00 | -1.53 | 0.14 |
| Control Low | Control Medium | 51.20 | 29.40 | 16.00 | 1.74 | 0.10 |
| Control Low | Control High | 19.40 | 29.40 | 16.00 | 0.66 | 0.52 |
| Control Low | Warm | -5.40 | 29.40 | 16.00 | -0.18 | 0.86 |
| Control Low | Warm Low | 3.53 | 29.40 | 16.00 | 0.12 | 0.91 |
| Control Low | Warm Medium | -17.87 | 29.40 | 16.00 | -0.61 | 0.55 |
| Control Low | Warm High | 26.47 | 29.40 | 16.00 | 0.90 | 0.38 |
| Control Medium | Control High | -31.80 | 29.40 | 16.00 | -1.08 | 0.30 |
| Control Medium | Warm | -56.60 | 29.40 | 16.00 | -1.92 | 0.07 |
| Control Medium | Warm Low | -47.67 | 29.40 | 16.00 | -1.62 | 0.12 |
| Control Medium | Warm Medium | -69.07 | 29.40 | 16.00 | -2.35 | 0.03 |
| Control Medium | Warm High | -24.73 | 29.40 | 16.00 | -0.84 | 0.41 |
| Control High | Warm | -24.80 | 29.40 | 16.00 | -0.84 | 0.41 |
| Control High | Warm Low | -15.87 | 29.40 | 16.00 | -0.54 | 0.60 |
| Control High | Warm Medium | -37.27 | 29.40 | 16.00 | -1.27 | 0.22 |
| Control High | Warm High | 7.07 | 29.40 | 16.00 | 0.24 | 0.81 |
| Warm | Warm Low | 8.93 | 29.40 | 16.00 | 0.30 | 0.77 |
| Warm | Warm Medium | -12.47 | 29.40 | 16.00 | -0.42 | 0.68 |
| Warm | Warm High | 31.87 | 29.40 | 16.00 | 1.08 | 0.29 |
| Warm Low | Warm Medium | -21.40 | 29.40 | 16.00 | -0.73 | 0.48 |
| Warm Low | Warm High | 22.93 | 29.40 | 16.00 | 0.78 | 0.45 |
| Warm Medium | Warm High | 44.33 | 29.40 | 16.00 | 1.51 | 0.15 |

Supplementary Table 8. GAM statistical summary for Black White Test crosses to the white zone for temperature (15°C and 20°C) and microplastic (0, 1, 10, 100 mg/g) treatments (n = 15).

| Fixed Effect | Estimate | SE | DF | t Value | P Value |
| --- | --- | --- | --- | --- | --- |
| Control Temperature | 2.03 | 0.15 | 96.00 | 13.35 | 0 |
| Warm Temperature | 0.51 | 0.14 | 19.00 | 3.76 | 0.0013 |
| Plastic Low | -0.32 | 0.19 | 19.00 | -1.68 | 0.11 |
| Plastic Medium | 0.07 | 0.19 | 19.00 | 0.39 | 0.70 |
| Plastic High | -0.27 | 0.19 | 19.00 | -1.42 | 0.17 |

Supplementary Table 9. GLMER statistical summary for Black White Test frequency of visits to the white zone used in the OR calculations for temperature (15°C and 20°C) and microplastic (0, 1, 10, 100 mg/g) treatments (n = 15).

| Fixed Effect | Estimate | SE | Z Value | P Value |
| --- | --- | --- | --- | --- |
| Control Temperature | 0.94 | 0.63 | 1.49 | 0.14 |
| Warm Temperature | 1.56 | 0.58 | 2.69 | 0.01 |
| Plastic Low | -1.35 | 0.80 | -1.70 | 0.09 |
| Plastic Medium | -0.06 | 0.84 | -0.07 | 0.95 |
| Plastic High | -1.05 | 0.80 | -1.31 | 0.19 |

Supplementary Table 10. LMER statistical summary for individual change in length for temperature (15°C and 20°C) and microplastic (0, 1, 10, 100 mg/g) treatments (n = 15).

| Fixed Effect | Estimate | SE | DF | t Value | P Value |
| --- | --- | --- | --- | --- | --- |
| Control Temperature | 1.80 | 1.40 | 115.00 | 1.29 | 0.20 |
| Warm Temperature | -0.40 | 1.25 | 115.00 | -0.32 | 0.75 |
| Plastic Low | 0.61 | 1.77 | 115.00 | 0.35 | 0.73 |
| Plastic Medium | 2.25 | 1.77 | 115.00 | 1.27 | 0.21 |
| Plastic High | 0.94 | 1.77 | 115.00 | 0.53 | 0.60 |

Supplementary Table 11. LMER statistical summary for individual change in weight for temperature (15°C and 20°C) and microplastic (0, 1, 10, 100 mg/g) treatments (n = 15).

| Fixed Effect | Estimate | SE | DF | t Value | P Value |
| --- | --- | --- | --- | --- | --- |
| Control Temperature | 0.01 | 0.11 | 115.00 | 0.12 | 0.91 |
| Warm Temperature | -0.13 | 0.10 | 115.00 | -1.32 | 0.19 |
| Plastic Low | 0.22 | 0.14 | 115.00 | 1.56 | 0.12 |
| Plastic Medium | 0.22 | 0.14 | 115.00 | 1.55 | 0.12 |
| Plastic High | 0.23 | 0.14 | 115.00 | 1.59 | 0.12 |

Supplementary Table 12. LMER statistical summary for Condition Factor for temperature (15°C and 20°C) and microplastic (0, 1, 10, 100 mg/g) treatments (n = 15).

| Fixed Effect | Estimate | SE | DF | t Value | P Value |
| --- | --- | --- | --- | --- | --- |
| Control Temperature | 0.90 | 0.01 | 115.00 | 65.72 | <2E-16 |
| Warm Temperature | -0.02 | 0.01 | 115.00 | -1.71 | 0.09 |
| Plastic Low | 0.01 | 0.02 | 115.00 | 0.69 | 0.49 |
| Plastic Medium | 0.01 | 0.02 | 115.00 | 0.52 | 0.61 |
| Plastic High | 0.04 | 0.02 | 115.00 | 2.39 | 0.02 |

Supplementary Table 13. Sidak comparisons of condition factor for temperature (15°C and 20°C) and microplastic (0, 1, 10, 100 mg/g) treatments (n = 15).

| Sidak Pairwise Comparison Treatment 1 | Sidak Pairwise Comparison Treatment 2 | Estimate | SE | DF | t Value | P Value |
| --- | --- | --- | --- | --- | --- | --- |
| Control | Control Low | -0.012 | 0.017 | 19 | -0.69 | 1.00 |
| Control | Control Medium | -0.009 | 0.017 | 19 | -0.52 | 1.00 |
| Control | Control High | -0.042 | 0.017 | 19 | -2.39 | 0.30 |
| Control | Warm | 0.021 | 0.012 | 19 | 1.71 | 0.68 |
| Control | Warm Low | 0.009 | 0.021 | 19 | 0.42 | 1.00 |
| Control | Warm Medium | 0.012 | 0.021 | 19 | 0.57 | 1.00 |
| Control | Warm High | -0.021 | 0.021 | 19 | -0.96 | 0.97 |
| Control Low | Control Medium | 0.003 | 0.017 | 19 | 0.18 | 1.00 |
| Control Low | Control High | -0.029 | 0.017 | 19 | -1.69 | 0.69 |
| Control Low | Warm | 0.033 | 0.021 | 19 | 1.55 | 0.77 |
| Control Low | Warm Low | 0.021 | 0.012 | 19 | 1.71 | 0.68 |
| Control Low | Warm Medium | 0.024 | 0.021 | 19 | 1.13 | 0.94 |
| Control Low | Warm High | -0.008 | 0.021 | 19 | -0.40 | 1.00 |
| Control Medium | Control High | -0.033 | 0.017 | 19 | -1.87 | 0.58 |
| Control Medium | Warm | 0.030 | 0.021 | 19 | 1.41 | 0.84 |
| Control Medium | Warm Low | 0.018 | 0.021 | 19 | 0.84 | 0.99 |
| Control Medium | Warm Medium | 0.021 | 0.012 | 19 | 1.71 | 0.68 |
| Control Medium | Warm High | -0.012 | 0.021 | 19 | -0.54 | 1.00 |
| Control High | Warm | 0.063 | 0.021 | 19 | 2.94 | 0.12 |
| Control High | Warm Low | 0.051 | 0.021 | 19 | 2.37 | 0.31 |
| Control High | Warm Medium | 0.054 | 0.021 | 19 | 2.51 | 0.25 |
| Control High | Warm High | 0.021 | 0.012 | 19 | 1.71 | 0.68 |
| Warm | Warm Low | -0.012 | 0.017 | 19 | -0.69 | 1.00 |
| Warm | Warm Medium | -0.009 | 0.017 | 19 | -0.52 | 1.00 |
| Warm | Warm High | -0.042 | 0.017 | 19 | -2.39 | 0.30 |
| Warm Low | Warm Medium | 0.003 | 0.017 | 19 | 0.18 | 1.00 |
| Warm Low | Warm High | -0.029 | 0.017 | 19 | -1.69 | 0.69 |
| Warm Medium | Warm High | -0.033 | 0.017 | 19 | -1.87 | 0.58 |

Supplementary Table 14. LMER statistical summary of Critical Thermal Maximum for temperature (15°C and 20°C) and microplastic (0, 1, 10, 100 mg/g) treatments (n = 11-12). Temperature and Plastic as independent fixed effects and tanks as nested factor.

| Fixed Effect | Estimate | SE | DF | t-value | P value |
| --- | --- | --- | --- | --- | --- |
| Control Temperature | 32.72 | 0.10 | 18.94 | 332.67 | 2.00E-16 |
| Warm Temperature | 1.96 | 0.09 | 19.09 | 22.28 | 3.99E-15 |
| Plastic Low | -0.45 | 0.12 | 18.90 | -3.62 | 0.0018 |
| Plastic Medium | -0.74 | 0.12 | 18.90 | -5.93 | 1.06E-05 |
| Plastic High | -0.36 | 0.13 | 19.28 | -2.87 | 0.0097 |

Supplementary Table 15. Estimated marginal mean tank Critical Thermal Maximum summary table for temperature (15°C and 20°C) and microplastic (0, 1, 10, 100 mg/g) treatments (n = 11-12).

| Treatment | EMM CTmax (°C) | SE | Lower CI | Upper CI | Degrees Freedom |
| --- | --- | --- | --- | --- | --- |
| Control | 32.7 | 0.1 | 32.5 | 32.9 | 18.8 |
| Control Low | 32.3 | 0.1 | 32.1 | 32.5 | 18.8 |
| Control Medium | 32.0 | 0.1 | 31.8 | 32.2 | 18.8 |
| Control High | 32.4 | 0.1 | 32.2 | 32.6 | 19.1 |
| Warm | 34.7 | 0.1 | 34.5 | 34.9 | 18.8 |
| Warm Low | 34.2 | 0.1 | 34.0 | 34.4 | 18.8 |
| Warm Medium | 33.9 | 0.1 | 33.7 | 34.2 | 18.8 |
| Warm High | 34.3 | 0.1 | 34.1 | 34.5 | 19.7 |

Supplementary Table 16. Sidak comparisons of Critical Thermal Maximum values for temperature (15°C and 20°C) and microplastic (0, 1, 10, 100 mg/g) treatments (n = 11-12).

| Sidak Pairwise Comparison Treatment 1 | Sidak Pairwise Comparison Treatment 2 | Estimate | SE | DF | t Value | P Value |
| --- | --- | --- | --- | --- | --- | --- |
| Control | Control Low | 0.45 | 0.12 | 18.80 | 3.62 | 0.03 |
| Control | Control Medium | 0.74 | 0.12 | 18.80 | 5.93 | 0.00 |
| Control | Control High | 0.36 | 0.13 | 19.20 | 2.87 | 0.14 |
| Control | Warm | -1.96 | 0.09 | 19.00 | -22.28 | <.0001 |
| Control | Warm Low | -1.51 | 0.15 | 18.90 | -9.94 | <.0001 |
| Control | Warm Medium | -1.23 | 0.15 | 18.90 | -8.05 | <.0001 |
| Control | Warm High | -1.61 | 0.15 | 19.40 | -10.46 | <.0001 |
| Control Low | Control Medium | 0.29 | 0.12 | 18.80 | 2.31 | 0.34 |
| Control Low | Control High | -0.09 | 0.13 | 19.20 | -0.73 | 0.99 |
| Control Low | Warm | -2.41 | 0.15 | 18.90 | -15.84 | <.0001 |
| Control Low | Warm Low | -1.96 | 0.09 | 19.00 | -22.28 | <.0001 |
| Control Low | Warm Medium | -1.68 | 0.15 | 18.90 | -11.00 | <.0001 |
| Control Low | Warm High | -2.06 | 0.15 | 19.40 | -13.39 | <.0001 |
| Control Medium | Control High | -0.38 | 0.13 | 19.20 | -3.03 | 0.10 |
| Control Medium | Warm | -2.70 | 0.15 | 18.90 | -17.73 | <.0001 |
| Control Medium | Warm Low | -2.25 | 0.15 | 18.90 | -14.77 | <.0001 |
| Control Medium | Warm Medium | -1.96 | 0.09 | 19.00 | -22.28 | <.0001 |
| Control Medium | Warm High | -2.34 | 0.15 | 19.40 | -15.26 | <.0001 |
| Control High | Warm | -2.32 | 0.15 | 18.90 | -15.24 | <.0001 |
| Control High | Warm Low | -1.87 | 0.15 | 18.90 | -12.29 | <.0001 |
| Control High | Warm Medium | -1.59 | 0.15 | 18.90 | -10.40 | <.0001 |
| Control High | Warm High | -1.96 | 0.09 | 19.00 | -22.28 | <.0001 |
| Warm | Warm Low | 0.45 | 0.12 | 18.80 | 3.62 | 0.03 |
| Warm | Warm Medium | 0.74 | 0.12 | 18.80 | 5.93 | 0.00 |
| Warm | Warm High | 0.36 | 0.13 | 19.20 | 2.87 | 0.14 |
| Warm Low | Warm Medium | 0.29 | 0.12 | 18.80 | 2.31 | 0.34 |
| Warm Low | Warm High | -0.09 | 0.13 | 19.20 | -0.73 | 0.99 |
| Warm Medium | Warm High | -0.38 | 0.13 | 19.20 | -3.03 | 0.10 |
